# PRC2-mediated repression is essential to maintain identity and function of differentiated dopaminergic and serotonergic neurons

**DOI:** 10.1101/2022.01.24.477226

**Authors:** Konstantinos Toskas, Behzad Yaghmaeian-Salmani, Olga Skiteva, Wojciech Paslawski, Linda Gillberg, Vasiliki Skara, Irene Antoniou, Erik Södersten, Per Svenningsson, Karima Chergui, Markus Ringnér, Thomas Perlmann, Johan Holmberg

## Abstract

How neurons in the CNS can maintain cellular identity over an entire lifespan remains largely unknown. Here we show that long-term maintenance of identity in differentiated dopaminergic and serotonergic neurons is critically reliant on the Polycomb repressive complex 2 (PRC2). Deletion of the obligate PRC2-component, *Eed*, in these neurons, resulted in global loss of H3K27me3, followed by a gradual activation of genes harbouring both H3K27me3 and H3K9me3 modifications. Notably, H3K9me3 was also lost at these PRC2-targets prior to gene activation. Neuronal survival was not compromised, instead there was a reduction in subtype specific gene expression as well as a progressive impairment of dopaminergic or serotonergic neuronal function leading to behavioural deficits characteristic of Parkinson’s disease (PD) or mood disorders, respectively. Single cell analysis revealed an unexpected subtype specific vulnerability to loss of PRC2-repression in dopamine neurons of the substantia nigra, the neurons primarily affected in PD. Taken together, our study reveals that a PRC2-dependent non-permissive chromatin state is essential to maintain subtype identity and function of dopaminergic and serotonergic neurons.

## Introduction

The brain contains a large number of different neuronal subtypes that maintain their distinct cellular identities over several decades despite continuous environmental fluctuation. Apart from the instructive information provided by transcription factors controlling cell type-specific gene programs, there is also a need to maintain silencing of transcriptional programs governing other cellular fates (*1, 2*). The mechanisms governing permanent repression of aberrant transcription in mature neurons are not well understood. Within this context, it is key to understand mechanisms regulating chromatin structure, e.g., dynamic modification of histones and how this is coupled to changes in gene expression. A prominent example of how chromatin associated gene silencing contributes to maintained cellular identity, is the sustained repression of *Hox* genes during segmentation of the *Drosophila melanogaster* embryo, which is dependent on polycomb group proteins (*3*). The Polycomb repressive complex 2 (PRC2) maintains established cell-type-specific gene repression through the deposition of the repressive histone modification H3K27me3 in promoter regions of silenced genes, thus facilitating chromatin compaction (*4–6*). PRC2 is required for proper differentiation during the development of the vertebrate CNS (*7, 8*), but whether PRC2 is an essential component for maintaining differentiated neuronal identity remains unclear. H3K27me3 is also found in domains which harbour the H3K4me3 modification associated with active transcription and these bivalent domains have been proposed to be silent but “poised” for rapid activation at a later developmental stage (*9, 10*). Thus, H3K4me3/H3K27me3 bivalency potentially represents a more relaxed chromatin state, amenable to rapid activation.

Efforts have been directed to understand how deviant gene regulation is involved in neurodegenerative and psychiatric disorders (*11*). The complex aetiology, often lacking a distinct and identifiable genetic component, of many of such pathological conditions suggests that alterations of the epigenome contributes to the disease (*11*). Changes in PRC2-activity and in H3K27me3 levels and distribution have been associated with neurodegenerative disease (*12–14*) and mood disorders (*15*), however, if or how these processes contribute to disease remains poorly understood.

To understand fundamental molecular mechanisms underlying the role of gene repression for maintenance of neuronal identity and function, we have focused on the well-defined mDA- and 5HT- neuronal subpopulations as model systems. These monoaminergic neurons are involved in several psychiatric disorders and drug addiction. In addition, degeneration of mDA neurons in the Substantia nigra *pars compacta* (SN*pc*) is a hallmark of Parkinson’s disease (PD) and dysregulated serotonergic function is causing depression, anxiety and contributes to L-DOPA dyskinesia in PD-patients (*16*). Hence, it is also relevant from a clinical perspective to understand basal mechanisms important for maintaining intact mDA and 5HT neuronal identity and function. Notably, in a preclinical model of PD, mDA neurons treated with the neurotoxin 6-OHDA exhibited substantial reduction in H3K27me3 levels (*17*). Besides the well-known toxic effects of 6-OHDA on mitochondria resulting in increased free radicals, this also couples exposure to PD-associated cellular stressors to the induction of a more relaxed chromatin state and potential de-repression of aberrant non-mDA genes. Furthermore, exposure to L-DOPA in a mouse model of PD leads to loss of Polycomb mediated repression (*12*). In differentiated medium spiny neurons, it has been shown that loss of PRC2 activity induced expression of cell death promoting genes which resulted in neuronal loss, leading to neurodegeneration (*6*). In these neurons de-repression primarily affected bivalent “poised” H3K4me3/H3K27me3 genes.

H3K9me3 is an additional histone modification involved in the establishment and maintenance of heterochromatin (*18*). Primarily associated with constitutive heterochromatin at centromeres and telomeres, H3K9me3 has also been shown to regulate facultative heterochromatin, provide an obstacle for cell reprogramming, and be required for establishment and maintenance of cellular identity (*18, 19*). Furthermore, H3K9me3 has been shown to be disrupted along with H3K27me3 upon 6-OHDA treatment (*17*). In a recent study we generated global integrated maps of transcription and histone modifications (H3K4me3, H3K27me3 and H3K9me3) of transitory as well as stable cellular states in mDA and 5HT neurons as well as their progenitors (*20*). This study also showed that in a mouse model of PD there was a significant enrichment of H3K27me3 targets among the upregulated genes implying a role for PRC2 in the transcriptional response to PD-associated cellular stressors.

To address the functional role of polycomb-mediated gene silencing for post-mitotic mDA- and 5HT- neuronal identity we have conditionally deleted the obligate PRC2-member *Eed*, which is necessary for PRC2 binding to H3K27me3 (*21*) and subsequent propagation of the modification, in both these neuronal subtypes. Taken together our study reveals a common logic in mDA- and 5HT-neurons wherein PRC2-activity is required for maintenance of subtype specific gene patterns and neuronal function, consequently loss of PRC2-function generates phenotypes which mirror key aspects of PD and mood disorders, without compromising neuronal survival. In addition, our single cell analysis reveals a specific vulnerability in mDA neurons of the SNpc to reduced H3K27me3 levels.

## Results

### Conditional deletion of *Eed* in differentiated mDA neurons

To address the role of PRC2 mediated repression in differentiated mDA neurons we generated a compound mouse mutant by crossing mice carrying floxed alleles for a ribosomal protein fused to mCherry (RPL10a-mCherry). The *Rpl10a-mCherry^flox/flox^* mouse line was crossed with mice carrying a floxed *Eed* allele (*22*) and finally with *DatCre* (*23*) (Fig. 1A). This enabled deletion of *Eed* and expression of *mCherry* in post-mitotic mDA neurons under the control of *Dat* (*Slc6a3)* expression, which is first detected in midbrain at around embryonic day 13.5 (E13.5). In this study, the full compound mutants will be denominated *DatCreEed^fl/fl^.* Pups from the mutants were born at expected mendelian rations and were indistinguishable from wild type littermates. Immunostaining of sections showed that the mCHERRY reporter colocalized with the rate limiting enzyme for dopamine synthesis, Tyrosine Hydroxylase (TH) in both the VTA and SN*pc* of both wild-type and mutant mice (Fig. 1B-E).

**Figure 1.**
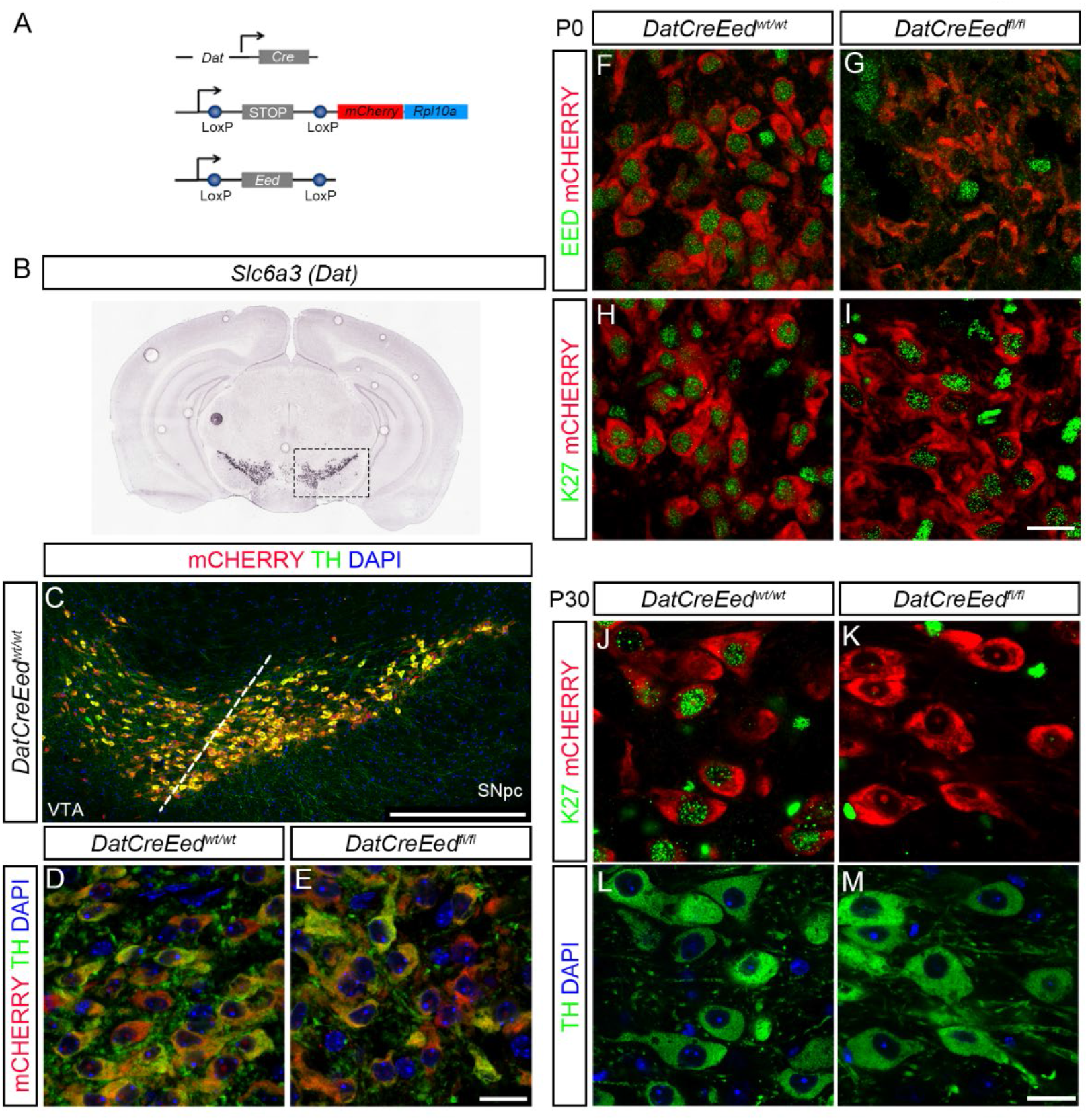
Deletion of *Eed* leads to progressive loss of H3K27me3 in differentiated mDA neurons. (**A**). Schematic representation of the three constructs used to generate the mutant mice. (**B**). *In situ* hybridization for *Slc6a3* (*Dat*) taken from the Allen Brain Atlas (mouse.brain-map.org, image credit: Allen Institute). (**C**). Immunostaining of ventral midbrain in *DatCreEed^wt/wt^* as indicated by box in **B,** showing overlap between mCHERRY and TH. (**D-E**). Immunostaining at 63x magnification showing overlap of TH and mCHERRY in SNpc of *DatCreEed^wt/wt^*-mice (**D**) and *DatCreEed^fl/f^*-mice (**E**). (**F**). mDA neurons double positive for EED and mCHERRY immunostaining at P0 in *DatCreEed^wt/wt^* SNpc. (**G**). Loss of EED in mCHERRY^+^-cells at P0 in *DatCreEed^ft/fl^* SNpc. (**H**). mDA neurons double positive for H3K27me3 and mCHERRY immunostaining at P0 in *DatCreEed^wt/wt^* SNpc. (**i**). mDA neurons double positive for H3K27me3 and mCHERRY immunostaining at P0 in *DatCreEed^fl/fl^* SNpc. (**J**). mDA neurons double positive for H3K27me3 and mCHERRY immunostaining at P30 in *DatCreEed^wt/wt^* SNpc. (**K**). Loss of H3K27me3 in mCHERRY^+^-cells at P30 in *DatCreEed^fl/fl^* SNpc. (**L**). Staining with TH and DAPI of the *DatCreEed^wt/wt^*-cells from panel **J**. (**M**). Staining with TH and DAPI of the *DatCreEed^fl/fl^*-cells from panel **K**. Scale bars in E, L:20mm

### Intact PRC2 function is required for long term maintenance of H3K27me3

To understand if induction of *Cre* resulted in deletion of EED protein at birth we performed immunostaining of midbrains from newborn pups at day 0 (P0) with an antibody specific for EED. In *DatCreEed^wt/wt^* mice there was clear nuclear EED immunoreactivity in mCHERRY^+^ mDA neurons (Fig. 1F) whereas mCHERRY^+^ cells in the *DatCreEed^fl/fl^* mutants exhibited lack of nuclear EED (Fig. 1G). To investigate whether the absence of EED protein in the mutants also resulted in loss of H3K27me3 we performed immunostaining with a H3K27me3 specific antibody. Although EED was lost in *DatCreEed^fl/fl^* H3K27me3 was retained (Fig. 1H-I), showing that the H3K27me3 modification is remarkably stable in post-mitotic mDA neurons. To further gauge if long term H3K27me3 stability depends on intact PRC2 function we stained midbrain sections of juvenile mice, at P30, which revealed a virtually complete lack of H3K27me3 immunoreactivity in *DatCreEed^fl/fl^*-mice (Fig. 1J-K). Notably, immunoreactivity levels of TH were retained in both genotypes (Fig. 1L-M).

### Progressive activation of silent non-mDA PRC2 targets and repression of mDA identity genes

To investigate the effects on global distribution of H3K27me3 and possible consequences of altered H3K27me3 levels for gene expression, we dissected out and prepared nuclei from dissected midbrains and used fluorescent activated cell sorting (FACS) to isolate mCHERRY^+^ nuclei from *DatCreEed^wt/wt^* and *DatCreEed^fl/fl^*-mice at four and eight months of age. Sorted nuclei were collected in batches of 1000. Batches were used to generate chromatin immunoprecipitation (ChIP) libraries for H3K27me3 (K27) but also for the permissive modification H3K4me3 (K4) and the facultative heterochromatin associated modification H3K9me3 (K9). In addition, one batch per mouse brain was utilized to generate libraries for bulk RNAseq.

Initially we examined the presence of K27 in the promoter region ± 10 kb around transcription start sites (TSS) together with expression levels in wild type cells which revealed an inverse correlation (Fig. 2A). We then determined how distinct chromatin states in the same promoter region correlated with expression levels in wild type cells at four months as described in our previous study (*20*). The results show that among the eight possible different states, the genes containing K4 exhibit the highest expression levels. Conversely, chromatin states harbouring K27 and/or K9 without K4 exhibit the lowest levels of expression (Supplementary Fig. S1A).

**Figure 2.**
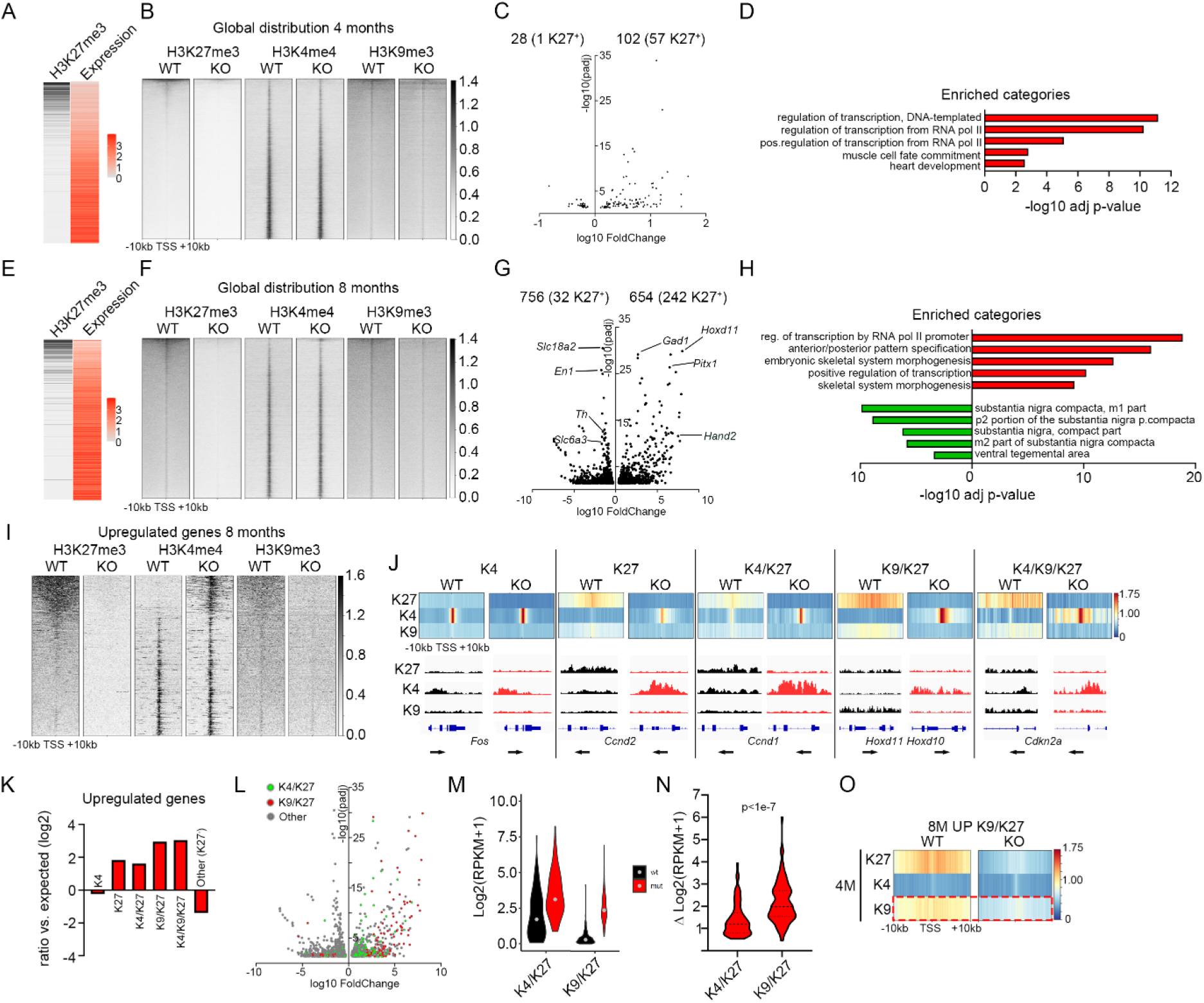
Loss of *Eed* results in progressive upregulation of PRC2 targets and reduced expression of mDA-neuronal genes. (**A**). Heatmaps showing inverse correlation between H3K27me3 (black) and expression levels (red). (**B**). Heat maps showing genome wide abundance of H3K27me3, H3K4me3 and H3K9me3 at 4 months in *DatCreEed^wt/wt^* (WT) and *DatCreEed^fl/fl^* (KO) at 10kb upstream and downstream of TSS at individual genes ranked by H3K27me3 abundance in the WT. (**C**). Volcano plot showing differentially regulated genes at 4 months in isolated mCHERRY^+^-nuclei from *DatCreEed^fl/fl^* ventral midbrain. Number of upregulated and downregulated genes are indicated above the plot, with the number of H3K27me3^+^-genes within brackets. (**D**). Enriched categories for upregulated genes (red) (GO biological process) are characterized by activation of transcription and early developmental non- neuronal processes. (**E**). Heatmaps showing inverse correlation between H3K27me3 (black) and expression levels (red). (**F**). Heat maps showing genome wide abundance of H3K27me3, H3K4me3 and H3K9me3 at 8 months in *DatCreEed^wt/wt^* (WT) and *DatCreEed^fl/fl^* (KO) at 10kb upstream and downstream of TSS at individual genes ranked by H3K27me3 abundance in the WT. (**G**). Volcano plot showing differentially regulated genes in isolated mCHERRY^+^-nuclei from *DatCreEed^fl/fl^* ventral midbrain. Number of upregulated and downregulated genes are indicated above the plot, with the number of H3K27me3^+^-genes within brackets. Examples of upregulated genes that are PRC2 targets in *DatCreEed^wt/wt^* mDA neurons are labelled on the right side of the plot. Examples of downregulated mDA-identity genes are labelled on the left side of the plot. (**H**). Enriched categories for upregulated genes (red) (GO biological process) are characterized by activation of transcription and early developmental non-neuronal processes, whereas downregulated genes (green) show enrichment for ventral midbrain categories (Up in Allen Brain Atlas, as calculated by Enrichr). (**I**). Heat maps showing abundance of H3K27me3, H3K4me3 and H3K9me3 at genes upregulated in 8 months *DatCreEed^fl/fl^* (KO) at 10kb upstream and downstream of TSS at individual genes ranked by H3K27me3 abundance in the WT. (**J**). Heatmap profiles of average H3K27me3, H3K4me3 and H3K9me3 RPKMs ±10kb around TSS of genes per defined chromatin states (denoted as K4, K27, K4/K27, K9/K27 and K4/K9/K27) in 8- month *DatCreEed^wt/wt^* (WT) and how these states are resolved in the *DatCreEed^fl/fl^* (KO) mDA-nuclei. Below each chromatin state, IGV-tracks exemplify how the chromatin states compare at representative genes between WT an KO. (**K**). Enrichment/depletion of chromatin states for upregulated genes in *DatCreEed^fl/fl^* mCHERRY^+^-cells at 8months. (**L**). Volcano plot as in **B** showing differentially regulated genes belonging to H3K4me3/H3K27me3 (green) or H3K9me3/H3K27me3 (red) chromatin states in WT cells. (**M**). Violin plot showing absolute expression level (log2(RPKM+1)) of upregulated genes in KO mDA-cells belonging to K4/K27 or K9/K27 chromatin states in 8-month WT mDA-cells. (**N**). Difference (Δ) in gene expression between WT and KO of genes belonging to K4/K27 or K9/K27. Student’s t-test. (**O**). Genes belonging to the K9/K27 chromatin state and upregulated in KO mDA-cells at 8-months have reduced K9 surrounding TSS already at 4 months despite no increased expression at 4 months.

We then investigated the consequences of *Eed* deletion for the distribution of K27, K4, and K9 at four and eight months. In four months mutant mDA neurons there was a substantial loss of K27, reducing the number of detected K27^+^ genes by 75.3% (2973 in wild type compared to 735 in mutants) (Fig. 2B, Supplementary table 1). To understand the consequence of such a major loss of K27 for gene expression, we integrated the expression data with the chromatin analysis. Despite the substantial global loss of K27, only 102 genes were significantly upregulated (adj p<0.05) (Fig. 2C, Supplementary table 2) and of those 57 genes were K27 targets (Fig. 2C, Supplementary Table 2), a 4.7-fold enrichment (Fisher’s exact test, p=5e-27) over the expected ratio. Out of the 57 upregulated K27 targets, 22 were determined as lacking K27 at ±10kb of TSS, in mutant nuclei. The 35 upregulated genes that still were determined as K27^+^, do retain K27 around the TSS, albeit at reduced levels (e.g., *Foxg1*, *Phox2b* and *Hand2*). Several quintessential PRC2 targets such as the *Hoxd* cluster, did not exhibit increased expression in the mutants. Despite a reduction in K27 enrichment in such genes, a significant proportion of the modification remained (Supplementary Fig. S1B), underscoring the resilience of this modification in mDA neurons lacking PRC2 activity. The upregulated genes exhibited strong enrichment of gene ontology (GO) categories associated with regulation of transcription and early developmental processes (GO Biological Process 2021, as calculated by Enrichr (*24, 25*)) (Fig. 2D). Concomitantly, loss of K27 also resulted in significant (adj p<0.05) downregulation of 28 genes, with one of them *(Myo7a)* being a K27 target (Fig. 2C).

ChIPseq and RNAseq analysis in nuclei sorted from 8 months old animals also showed an inverse correlation between K27 and expression level in WT nuclei (Fig. 2E). Furthermore, it revealed a complete loss of genes determined as K27^+^ in the *DatCreEed^fl/fl^* mutants (Fig. 2F, Supplementary Table 1). This loss was accompanied by a minor but noticeable increase in K4 and loss of K9 (Fig. 2F). Inspection of differentially expressed genes showed that 654 genes were significantly upregulated (Fig. 2G), with 55 of them upregulated at 4 months and 242/654 carrying the K27 modification (Fig. 2G, Supplementary Table 2), representing a 4.3-fold enrichment over expected (Fisher’s exact test, p=5e-95). Upregulated PRC2-targets included several members of the *Hox*-family, transcription factors involved in determining other cell fates during development e.g., *Pitx1, Gata2 and Foxd3*, stem cell factors such as *Pax6*, genes typically expressed in other neuronal types, e.g., *Gad1-2* and cell cycle regulators including *Ccnd1-2* and *Cdkn2a* (Supplementary table 2). In contrast to the four-month mutant mDA neurons, several members of the *Hoxd* clusters were upregulated at 8 months. This was reflected by acquisition of H3K4me3 combined with loss of K27 as well as K9 (Supplementary Fig. S1C). Moreover, in the 8-month-old mutant mDA neurons there were 756 significantly downregulated genes, with 32 of them carrying the K27 modification (Fig. 2G, Supplementary table 2), representing a two-fold reduction compared to what would be expected by chance (Fisher’s exact test, p=7e-5). Among the downregulated genes several transcription factors critical for mDA neuronal function were present, e.g., *En1/2*, *Nr4a2* (*Nurr1*), *Lmo3, Pitx3* and *Pou3f2* (Supplementary table 2). Upregulated genes exhibited strong enrichment of GO-categories associated with regulation of transcription and early developmental processes, whereas downregulated genes were enriched for ventral midbrain categories (GO Biological Process 2021 and Allen Brain Atlas Up, as calculated by Enrichr) (Fig 2H).

### Combined H3K9me3/H3K27me3 is associated with higher probability of de-repression upon loss of PRC2 activity

Inspection of the H3K27me3, H3K4me3 and H3K4me9 modifications ±10 kb around the TSSs of the genes upregulated in the mutants, revealed a pronounced increase of K4 and loss of K9 (Fig. 2I). To further understand whether any specific chromatin state in wild type mDA neurons would predispose for de-repression, we inspected the distribution of K4, K27, K4/K27, K9/K27 and K4/K9/K27 in wild type mDA nuclei and correlated them with the differentially expressed genes. Among the upregulated genes there was a significant enrichment in all states containing K27 but not in the K4-only, where there rather was a depletion (Fig. 2J-K). Closer inspection revealed that the most enriched chromatin states of upregulated genes in the mutants were those that contained both HK27me3 and HK9me3 (K4/K9/K27 and K9/K27) in wild type cells. Compared to genes with a “poised”, bivalent K4/K27 state in the wild type, which had a 3.1x enrichment, the K9/K27 state exhibited 7.8x and the K4/K9/K27 state 8.3x enrichment (Fig. 2K). In addition, when we compared the relative fold-change and statistical significance of the differentially expressed genes (DEGs) in the 8-months mutants, there was a clear difference between K4/K27 and K9/K27 genes, with K9/K27 genes generally exhibiting higher fold- increase and lower adjusted p-values (Fig. 2L). This was also true for the 4-months mutants (Supplementary Fig. S1D). To understand whether this difference was a consequence of K9/K27 genes being derepressed from absolute expression levels close to or equal to zero, we investigated the absolute expression levels of K4/K27 and K9/K27 genes in 8 months old wild type and mutants. This revealed that the wild type expression levels of K9/K27 genes were substantially lower than those of K4/K27 genes (Fig. 2M). However, the difference in absolute increase was significantly larger in the K9/K27 genes (Fig. 2N). Thus, this analysis revealed that presence of the additional heterochromatin modification K9 actually increases both the probability to activate repressed K27 genes as well as the magnitude of increased expression. Notably, GO analysis showed that K9/K27 genes are enriched for categories typically including regulation of transcription and early developmental events such embryonic organ morphogenesis, whereas K4/K27 genes includes GO categories associated with regulation of extracellular matrix, neuronal differentiation and proliferation (Supplementary Fig. S1E). The stronger enrichment of K9/K27 genes over K4/K27 genes prompted us to inspect the chromatin states at 4 months for genes that were upregulated at 8 months but not at 4 months. This analysis revealed that reduction of K9 occurs prior to de-repression of expression (Fig. 2O). Hence, the early loss of K9 is not a mere consequence of activated transcription but rather coupled to the loss of K27 at the same TSS. We have previously shown that early developmental regulators already silent and harbouring K27 in neural stem cells, gain K9 in differentiated mDA neurons (*20*) suggesting that both the acquisition and maintenance of K9 at these TSSs are closely connected and potentially dependent on PRC2 activity.

### Progressive loss of TH, dopamine and dopamine-associated metabolites upon loss of PRC2 activity

To understand the phenotypic consequences of the progressive loss of K27 we stained for TH at 4 and 8 months, both in the midbrain VTA/SNpc and in the striatal target region. At 4 months there was no apparent difference between wild type and mutant mice, neither in VTA/SNpc nor in striatum (Fig. 3A-B, G-H), suggesting that in the mutants the establishment of this circuitry during development is not perturbed. Reflecting the reduction of *Th* expression in 8-months mutants (Fig. 2G), there was a slight reduction of TH immunoreactivity in mutant midbrain (Fig. 3D, J). However, in the dorsal striatum of *DatCreEed^fl/fl^* mutants TH was completely lost while low levels of TH could still be detected in the nucleus accumbens (Fig. 3C, I). To investigate whether this loss of TH progressed over time, we performed staining of 16-months old mutants. At this time point also mCHERRY^+^ cells in the SN*pc* displayed an almost complete loss of TH, whereas cells in the mutant VTA still exhibited substantial TH immunoreactivity (Fig. 3E-F, K-L). The loss of TH immunostaining in mutant mice at 8 months could be the result of a progressive loss of established projections from the SN/VTA to their striatal targets. To investigate this possibility, we performed intracranial injections targeting the SN/VTA with AAV-vectors (pCAG-FLEX-EGFP-WPRE virus (*26*)) to anterograde trace the projections from midbrain to striatum at 8 months. Three weeks post-injection we sacrificed the animals for analysis. Green fluorescent protein (GFP) was exclusively expressed from the mDA neurons expressing the recombinant protein CRE at the site of injection (Fig. 3N, 3P), as well as extending until their target area in the striatum (Fig. 3M, 3O). Despite, an almost total absence of TH immunoreactivity in mutant striatum there was a strong GFP signal in all mutants analysed (Fig. 3M-P). Hence, the projections from midbrain to the striatal target area are largely intact in the *DatCreEed^fl/fl^*-mutants. Since it has been reported that *Ezh1/2* deletion in medium spiny neurons caused severe neuronal loss through cell death (*6*), we analysed and counted the mCHERRY^+^ cells in representative areas of 8 months old midbrains but failed to detect any significant difference in the number of cells between *DatCreEed^wt/wt^* control mice and *DatCreEed^fl/fl^* mutants (Fig. 3Q-S). We next wanted to understand whether the reduced levels of TH affected levels of dopamine and associated metabolites. To investigate this, we dissected out midbrain and striatum and performed high performance liquid chromatography (HPLC) to measure metabolites of the dopamine synthesis pathway. In *DatCreEed^fl/fl^* mutants both regions displayed a significant reduction of dopamine, DOPAC and homovanillic acid (HVA), (Fig. 3T). The decrease was most prominent in the striatum, where mutant animals exhibited a more than fivefold reduction of all three metabolites compared with control animals (Fig. 3T).

**Figure 3.**
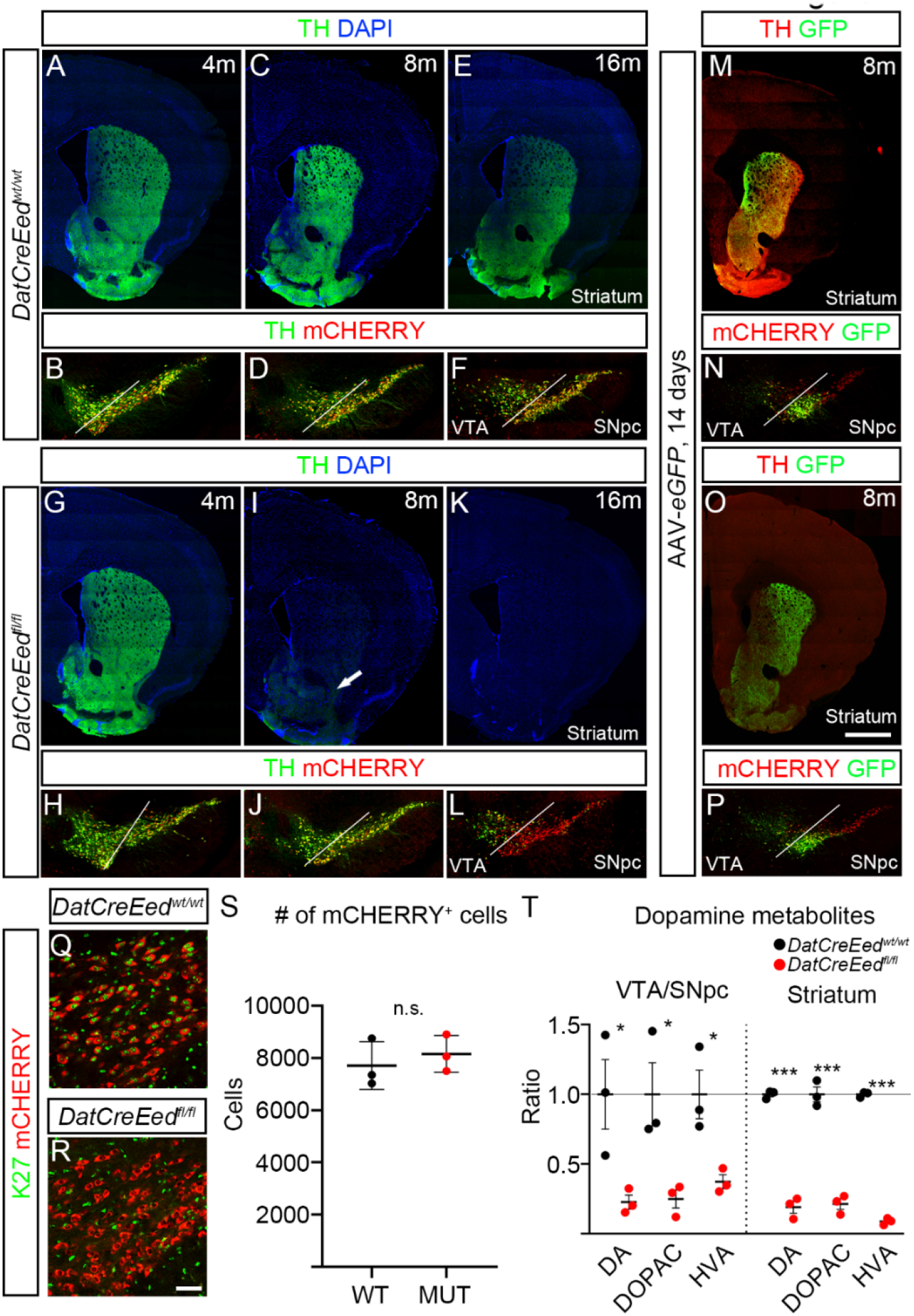
Reduced levels of TH and dopamine metabolites in striatum and midbrain upon inactivation of PRC2. (**A**). TH immunostaining in the striatum of *DatCreEed^wt/wt^*-mice at 4 months. (**B**). TH immunostaining and mCHERRY-fluorescence in the ventral midbrain of *DatCreEed^wt/wt^*-mice at 4 months. (**C**). TH immunostaining in the striatum of *DatCreEed^wt/wt^*-mice at 8 months. (**D**). TH immunostaining and mCHERRY-fluorescence in the ventral midbrain of *DatCreEed^wt/wt^*-mice at 8 months. (**E**). TH immunostaining in the striatum of *DatCreEed^wt/wt^*-mice at 16 months. (**F**). TH immunostaining and mCHERRY-fluorescence in the ventral midbrain of *DatCreEed^wt/wt^*-mice at 16 months. (**G**). TH immunostaining in the striatum of *DatCreEed^fl/fl^*-mice at 4 months. (**H**). TH immunostaining and mCHERRY-fluorescence in the ventral midbrain of *DatCreEed^fl/fl^*-mice at 4 months. (**I**). Reduced TH immunostaining in the striatum of *DatCreEed^fl/fl^*-mice at 8 months. Arrow indicates reduced albeit detectable TH immunostaining in the Nucleus Accumbens. (**J**). TH immunostaining and mCHERRY-fluorescence in the ventral midbrain of *DatCreEed^fl/fl^*-mice at 8 months. (**K**). Reduced TH immunostaining in the striatum of *DatCreEed^fl/fl^*-mice at 16 months. (**L**). TH immunostaining and mCHERRY-fluorescence in the ventral midbrain of *DatCreEed^fl/fl^*-mice at 16 months. (**M**). Expression of eGFP in striatum of *DatCreEed^wt/wt^*-mice 21days after injection of *AAV- eGFP* in ventral midbrain. (**N**). Site of injection of *AAV-eGFP* in *DatCreEed^wt/wt^*-mice 21 days post- injection. (**O**). Expression of eGFP in striatum of *DatCreEed^fl/fl^*-mice 21days after injection of *AAV-eGFP* in ventral midbrain. (**P**). Site of injection of *AAV-eGFP* in *DatCreEed^fl/fl^*-mice 21 days post-injection. (**Q**). H3K27me3 immunostaining in mCHERRY^+^-cells in SN of 8 months *DatCreEed^wt/wt^*-mice. (**R**). No H3K27me3 immunostaining in mCHERRY^+^-cells in SN of 8 months *DatCreEed^wt/wt^*-mice. (**S**). Quantification of mCHERRY^+^-cells in ventral midbrain of *DatCreEed^wt/wt^*-mice and *DatCreEed^fl/fl^*-mice at 8 months shows no loss of mCHERRY^+^-cells in the mutant midbrains. (**T**). Reduced levels of dopamine metabolites in the ventral midbrain and striatum of *DatCreEed^fl/fl^*-mice. In **T** *p<0.05, *** p<0.001, Unpaired t-test with Welch’s correction. Scale bar in O: 1000μm, in R: 50μm.

### Basic electrophysiological properties of mDA neurons are perturbed upon loss of PRC2 activity

The reduction in dopamine and associated metabolites combined with the loss of TH immunoreactivity in the striatal target area suggest that, besides loss of identity, a severe perturbation of mDA-neuron function also occurred in *DatCreEed^fl/fl^*-mutants. To address this, we analysed basal physiological properties through whole cell patch-clamp recordings in slice preparations from the 8- month mutant and wild type SNpc. Several of the measured parameters were significantly perturbed in the mutant midbrains (Fig. 4A-M). Cell capacitance was reduced (Fig. 4A) whereas membrane resistance was slightly increased (Fig. 4B). This implies that mutant mDA neurons exhibit smaller surface area and reduced open channel activity, generating a higher input resistance. Spontaneous pacemaker spiking typical of mDA neurons exhibited no difference in frequency (Fig. 4C), but the pacemaker pattern was disturbed with significant loss of consistency of interspike intervals (Fig. 4D-E). The hyperpolarization-activated current I_h_, mediated by hyperpolarization-activated cyclic nucleotide gated (HCN) channels, which contributes to mDA-neuronal pacemaker firing integrity, was reduced (Fig. 4F-G). Also, slow afterhyperpolarization current (AHC) generated by small-conductance Ca^2+^-sensitive K^+^ channels, was reduced (Fig. 4J-M). Even though the action potential amplitude was unchanged the threshold was decreased, whereas afterhyperpolarization (AHP) amplitude was decreased (Fig. 4I-L). These anomalies of spontaneous firing in the mDA-neuron population combined with the low levels of dopamine in the striatum indeed suggest severe loss-of mDA-neuronal function.

**Figure 4.**
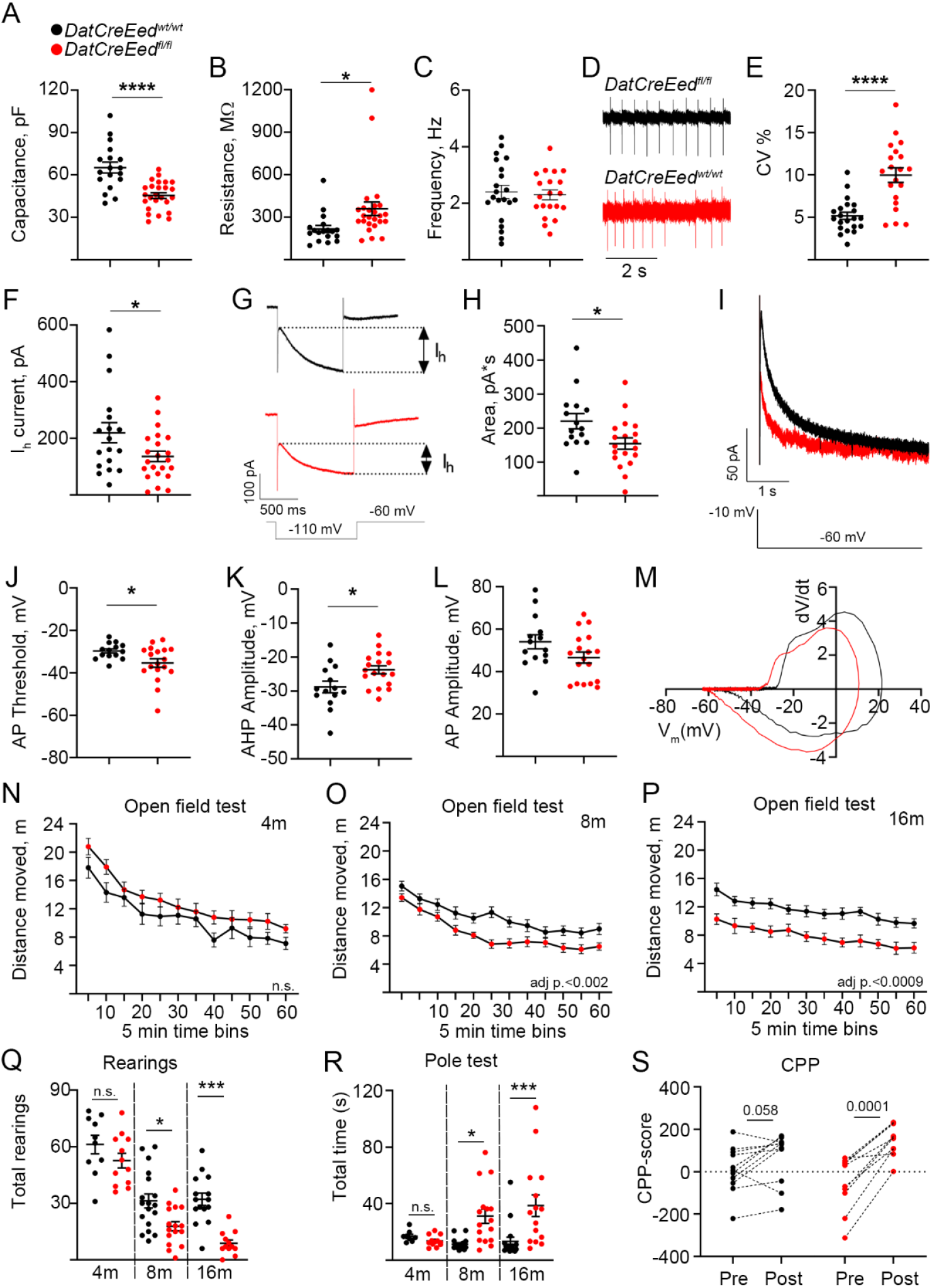
Altered electrophysiological properties and animal behaviour upon inactivation of PRC2. (**A**). Reduced capacitance in 8-month *DatCreEed^fl/fl^* mDA neurons. (**B**). Increased membrane resistance in 8-month *DatCreEed^fl/fl^* mDA neurons. (**C-E**). Increased coefficient of variation of interspike intervals of autonomous pacemaker currents in in 8-month *DatCreEed^fl/fl^* mDA neurons. (**F-G**). Decreased hyperpolarization current (I_h_) in 8-month *DatCreEed^fl/fl^* mDA neurons. (**H-I**). Decreased slow afterhyperpolarization current in 8-month *DatCreEed^fl/fl^* mDA neurons. (**J-K**). Action potential (AP) threshold is reduced (**J**), whereas afterhyperpolarization is decreased (**K**) in *DatCreEed^fl/fl^* mDA neurons. (**L**). AP amplitude was not significantly reduced in *DatCreEed^fl/fl^* mDA neurons. (**M**). Phase plot (dV/dt versus V_m_) of action potential in *DatCreEed^wt/wt^* and *DatCreEed^fl/fl^*-mDA neurons of the SNpc. (**N-P**). Open field test at 4 months (**N**), 8 months (**O**) and 16 months (**P**) shows progressive decrease in distance moved by *DatCreEed^fl/fl^*-mice. (**Q**). Progressive increase in total time needed for *DatCreEed^fl/fl^*-mice to complete the pole test. (**R**). Progressive decrease in number of rearings for *DatCreEed^fl/fl^*-mice. (**S**). CPP-score pre and post exposure to cocaine in wild type and mutant mice. In **A-M**, the data sets were checked for normality with Shapiro-Wilk test. For normally distributed data sets unpaired t-test was used (* - p < 0.05, **** - p < 0.0005). If the data sets did not pass the normality test – Mann-Whitney test was applied. In **N-P**, p-values calculated by Two-way repeated measures ANOVA. In **Q-R**, *p<0.05, ***p<0.001 calculated by One-way ANOVA with Tukey’s multiple comparisons test. In **S** p-values calculated with paired- t-test.

### Progressive impairment of overall locomotor activity and fine motor movement in *Eed* mutants

To test if reduced levels of TH and dopamine metabolites as well as altered electrophysiological properties would generate any behavioural consequences, we performed several behavioural tests to detect potential motor skill impairments in the mutant mice (*27*). Coordination, endurance and muscle strength were not affected in the mutants as shown by the rotarod and grip strength tests respectively. (Supplementary Fig. S2A-C). However, when analysed in the open field test, their overall locomotor activity exhibited progressive reduction with age as recorded from the total moved distance (Fig. 4M-O). In addition, the number of rearings was significantly reduced in the mutants (Fig. 4P). When challenged by the pole test, assessing fine locomotor function, mutant mice underscored a significant delay in initiation of descent as well as frequent failure to climb down the pole by falling sideways (Fig. 4Q). Hence, the mutants exhibited deficits primarily in the initiation of voluntarily movement as seen in the open field test, number of rearings and pole test. Whereas, when exposed to a forced movement paradigm testing coordination and balance, such as the rotarod, the mutant mice performed on par with the wild type mice. Motor function has mainly been coupled to the SNpc, whereas reward and motivation processes to a larger degree involve the VTA (*28*). Therefore, we challenged the mice by conditioning them to cocaine and examined whether perturbation in the reward system of 8-months old mutant mice occurred. We conducted conditional place preference (CPP) for cocaine, where a weak CPP occurred in the wild type mice (Fig. 4R). The CPP in mutant mice was more distinct (Fig. 4R), however the difference in post-test CPP between wild type and mutant mice was not significant, indicating that response to cocaine conditioning was unaffected by genotype. To investigate whether the motor response to cocaine was affected in the mutants we also performed an open field test after injection of 10mg/kg cocaine. In both wild type and mutant mice there was a substantial increase of movement compared to non-treated animals (Supplementary fig. S2D). However, there was no significant difference between cocaine-treated animals, with different genotypes (Supplementary fig. S2D). Thus, the capacity to respond to cocaine in the mutant is not impaired, implicating that this aspect of SNpc/VTA function is intact.

Taken together, the loss of *Eed* results in progressive loss of H3K27me3, leading to upregulation of Polycomb target genes and reduced expression of mDA neuronal genes. This loss of cellular identity severely disrupts mDA neuronal function at cellular level, ultimately altering the behaviour of *DatCreEed^fl/fl^*-mutants.

### Loss of PRC2-function in serotonergic neurons results in loss of cellular identity and function

An earlier study reported that cell-specific loss of PRC2 function in striatal medium spiny neurons (MSNs) caused substantial cell death (*6*), As mentioned, a corresponding cell death was not noted in the mDA neurons of *DatCreEed^fl/fl^*-mutants. To understand if deletion of *Eed* can cause loss of neuronal identity without cellular loss also in a different neuronal population we crossed the *Rpl10a- mCherry^flox/flox^/Eed^flox/flox^* mice with *SertCre-*mice(*29*). In this *SertCreEed^fl/fl^*-mutant, *Eed* is selectively deleted in hindbrain serotonergic neurons (5HT-neurons) expressing the serotonin transporter *Slc6a4* (a.k.a. *Sert.*) (Fig. 5A). Expression of *Slc6a4* first occurs in the hindbrain at around E12.5 of the mouse embryo. As in mDA neurons, there was in P0 mCHERRY^+^ 5HT-neurons a substantial reduction in EED immunoreactivity in the *SertCreEed^fl/fl^*-mutants, whereas levels of H3K27me3 remained as in *SertCreEed^wt/wt^* mice (Supplementary fig. S3A-D). At P40 the levels of H3K27me3 were not detectable in *SertCreEed^fl/fl^* mutant cells (Fig. 5B-C). To investigate possible consequences of *Eed* loss in 5HT- neurons we stained the hindbrains of wild type and mutants, with an antibody specific for the rate-limiting enzyme in serotonin synthesis, tryptophane hydroxylase 2 (TPH2). At four months of age the number of mCHERRY^+^ cells in the *SertCreEed^fl/fl^* mutants was unaltered compared to the wild type mice with a distinct overlap of TPH2 immunoreactivity and mCHERRY^+^ fluorescence in both wild type and mutant animals (Fig 5D, G). In contrast, at 8- and 16-months, immunostaining revealed a major loss of TPH2 in *SertCreEed^fl/fl^* mutants, but no loss of mCHERRY^+^-cells (Fig. 5E-I).

**Figure 5.**
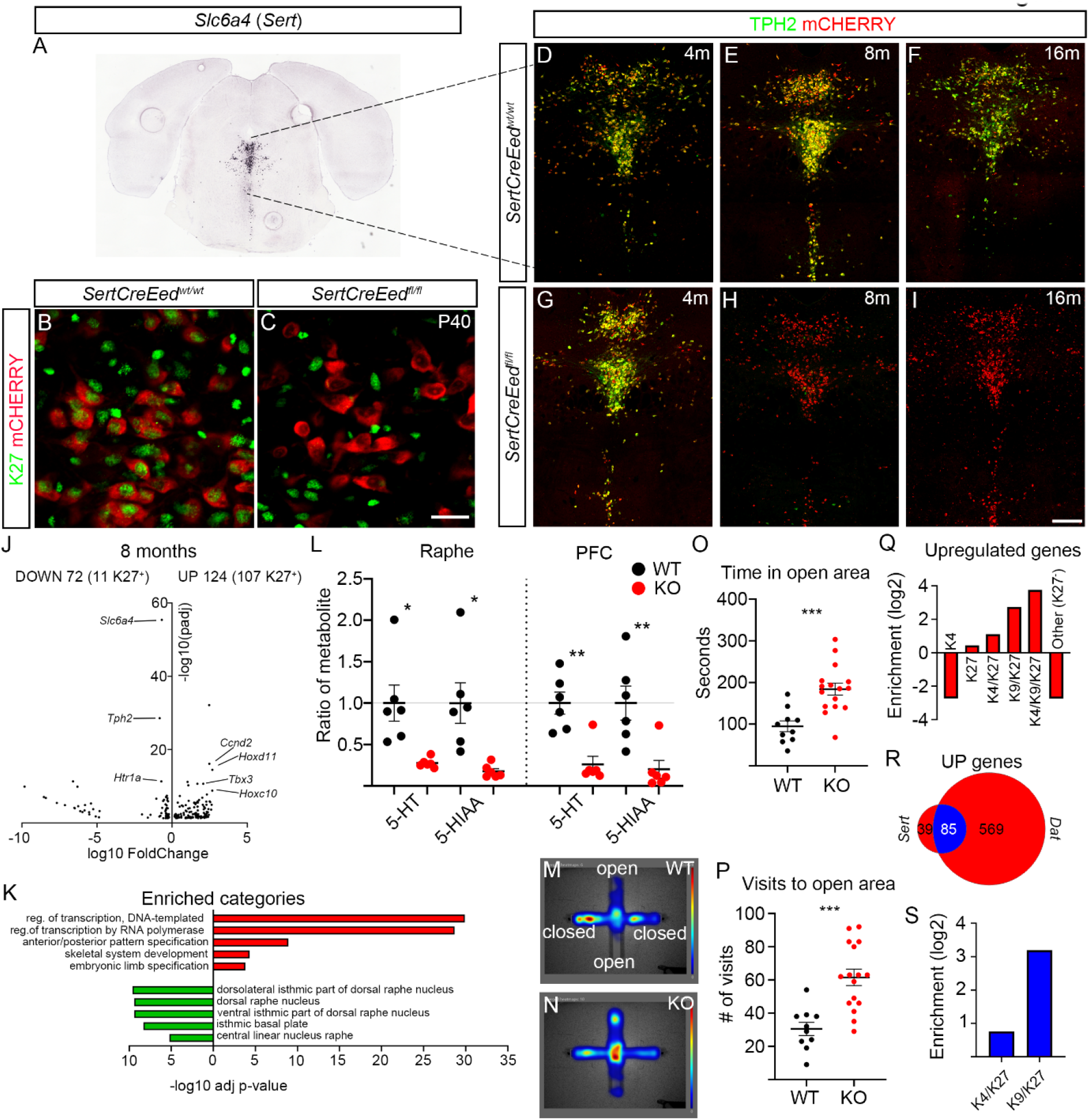
*Eed* deficiency in 5HT-neurons results in impaired 5HT-specific gene expression and function. (**A**). *In situ* hybridization for *Slc6a4* (*Sert*) taken from the Allen Brain Atlas (mouse.brain- map.org, image credit: Allen Institute). (**B**). Immunostaining of H3K27me3 in mCHERRY^+^-5HT-neurons in the dorsal raphe of *SertCreEed^wt/wt^*-mice. (**C**). Lack of H3K27me3 immunostaining in mCHERRY^+^-5HT- neurons in the dorsal raphe of *SertCreEed^fl/fl^*-mice. (**D-F**). TPH2 immunostaining localized in mCHERRY^+^ 5HT-neurons in the dorsal raphe of *SertCreEed^wt/wt^*-mice aged 4 months (**D**), 8 months (**E**) and 16 months (**F**). (**G-J**). Progressive loss of TPH2 in 5HT-neurons in the dorsal raphe of *SertCreEed^fl/fl^*-mice aged 4 months (**G**), 8months (**H**) and 16 months (**I**). (**J**). Volcano plot showing differentially regulated genes in isolated mCHERRY^+^-nuclei from *SertCreEed^fl/fl^* ventral midbrain. Number of upregulated and downregulated genes are indicated above the plot, with the number of H3K27me3^+^-genes within brackets. Examples of upregulated genes that are PRC2 targets in *SertCreEed^wt/wt^* mDA neurons are labelled on the right side of the plot. Examples of downregulated 5HT-identity genes are labelled on the left side of the plot. (**K**). Enriched categories for upregulated genes (red) (GO biological process) are characterized by activation of transcription and early developmental non-neuronal processes, whereas downregulated genes (green) show enrichment for raphe categories (Up in Allen Brain Atlas, as calculated in Enrichr). (**L**). Reduced levels of serotonin metabolites in the raphe and prefrontal cortex (PFC) of *SertCreEed^fl/fl^*-mice. (**M**). Heatmap of time spent in indicated areas in the elevated plus maze (EPM) for *SertCreEed^wt/wt^*-mice. (**N**). Heatmap of time spent in indicated areas in the elevated plus maze (EPM) for *SertCreEed^fl/fl^*-mice. (**O**). Increased time spent in open area for *SertCreEed^fl/fl^*-mice in the EPM. Unpaired t-test with Welch’s correction. (**P**). Increased number of visits to open area for *SertCreEed^fl/fl^*-mice in the EPM. Unpaired t-test with Welch’s correction. (**Q**). Enrichment of upregulated genes in *SertCreEed^fl/fl^* mCHERRY^+^ nuclei at the different chromatin states. (**R**). Overlap of upregulated genes in *DatCreEed^fl/fl^*-mice and *SertCreEed^fl/fl^*-mice, K27^+^ genes indicated in blue. (**S**). Commonly upregulated genes in *DatCreEed^fl/fl^*-mice and *SertCreEed^fl/fl^*-mice are more enriched for the K9K/K27 chromatin state than the K4/K27 state. In **L**, **O** and **P** *p<0.05, **p<0.01, *** p<0.001, Unpaired t-test with Welch’s correction. Scale bar in C: 20μm, in I: 200μm.

We proceeded to perform RNA-seq analysis of sorted 5HT-nuclei at 4 months which revealed upregulation of 84 transcripts of which 36 were K27^+^, as previously determined (*20*). Twenty-six transcripts were downregulated, of which two were K27^+^ (Supplementary Fig. S3E). Similar analysis of sorted 5HT-nuclei at 8 months revealed 124 upregulated genes of which 107 were K27^+^ (Fig. 5J). As in the mutant mDA nuclei the upregulated genes were enriched for several members of the *Hox*-family, transcription factors involved in determining other cell fates during development e.g., *Gata6*, *Foxg1* and *Dlx1*, stem cell factors such as *Pax6*, genes typically expressed in other neuronal types, e.g., *Gad1* and cell cycle regulators including *Ccnd2* and *Cdkn2a* (Fig. 5J and Supplementary Table 3). Downregulated genes numbered 72 and included 5HT-specific genes such as *Slc6a4, Tph2* and *Htr1a* (Fig. 5J and Supplementary Table 3). Of these 72 genes, 11 were H3K27me3^+^, which does not constitute an enrichment (Fig. 5J). GO analysis of the differentially expressed genes showed that among upregulated genes, there was a strong enrichment of categories related to transcriptional activation and early developmental processes. For the downregulated genes there was an enrichment of dorsal raphe nucleus associated categories (GO Biological Process 2021 and Allen Brain Atlas Up, as calculated by Enrichr) (Fig. 5K). The substantial reduction of TPH2 in 8-month mutants caused a decrease in levels of serotonin (5-HT) and the serotonin associated metabolite (5-HIAA) both in the hindbrain and in the prefrontal cortex target area as measured by HPLC (Fig. 5L).

To understand whether loss of serotonergic identity was followed by altered behaviour, as seen in the *DatCreEed^fl/fl^* mutants, we investigated whether aspects of behaviour that depend on an intact 5HT- function were perturbed in the 5HT-mutants. To examine if loss of *Eed* evoked depressive behaviour, we subjected 8-month *SertCreEed^wt/wt^* and *SertCreEed^fl/fl^*-mice to the forced swim test. Notably, there was no significant difference between wild-type and mutant mice for the time they spent swimming (Supplementary Fig. S3F). However, when subjected to the elevated plus maze (EPM) (*30*) the mutants spent significantly more time in the open arms and visited them more frequently (Fig 5M-P). This change in behaviour could be a consequence of hyperactivity or less anxiety, behavioural phenotypes which has previously been associated with deficient serotonin neurotransmission (*31, 32*).

### Activated PRC2 targets are enriched for H3K9me3 in 5HT-nuclei

The strong enrichment of K27 targets among the upregulated genes is reminiscent of the effects of *Eed*-deletion in mDA neurons. Since the presence of the heterochromatin modification H3K9me3 was associated with a higher probability of de-repression and activated transcription in mutant mDA nuclei we assigned upregulated genes in the *SertCreEed^fl/fl^* nuclei to the same chromatin states by utilizing the data set we generated for wild type 5HT-neurons in our previous study (*20*). This analysis showed that also in mutant 5HT nuclei the presence of K9 in a chromatin state was associated with a higher probability to activate transcription than the presence of K4 (Fig. 5Q).

### Common enrichment of upregulated H3K9me3/H3K27me3 targets in mutant mDA- and 5HT- neurons

Since we noted that transcriptional response to loss of *Eed* included similar PRC2 targets in mDA and 5HT neurons we compared the overlap of up-regulated genes at 8 months. Indeed, there was a large overlap of upregulated genes with 85 common transcripts out of 124 (5HT nuclei) and 654 (mDA nuclei) (Fig. 5R). This represents a more than 24-fold higher number than expected by chance (p<2.2e- 16, Fisher’s exact test). Of these 85 transcripts, 83 were K27^+^ PRC2 targets in 5HT neurons and 77 in mDA neurons, which represents a substantial enrichment in both cell types. In addition, the 85 commonly upregulated targets were also strongly enriched for the K9/K27-state when compared to the “poised” K4/K27-state (Fig. 5S).

When comparing Erichr (*24, 25*) analysis between the upregulated genes in mDA and 5HT neurons the similarity is clear. In both neuronal cell-types there was a strong enrichment of early developmental regulators, e.g., *Hox*-genes (Fig. 2G, 5J). Furthermore, another similarity is that both types of neurons exhibit reduced expression of transcripts specific to their identity, e.g., *Th, Slc6a3*, *En1, Nr4a2 and Pitx3* in the mDA neurons and *Tph2, Htr1a, Slc6a4* and *Htr5b* in 5HT-neurons (Fig. 2G, 5J). Notably, in both mutants substantially increased expression of genes previously described as “death promoting” (e.g., *Cdkn2a*, *Hoxa5, Wt1*) was evident (Supplementary Tables 2 & 3) but without inducing any cell death, which clearly distinguishes these neuronal subtypes from medium spiny neurons (*6*).

### Single nuclei expression analysis reveals SNpc specific vulnerability to loss of PRC2 activity

The selective and progressive loss of TH in the SNpc at 8 and 16 months (Fig. 3) indicates that different mDA-neuron subtypes may respond differently to PRC2 deficiency. To explore whether the changes in gene expression upon loss of H3K27me3 is distinct between mDA-neuron subgroups, gene expression was analysed by single nuclei RNA sequencing (snRNAseq) of sorted mCHERRY^+^ nuclei from the midbrain of 8-months old wild type and mutant mice. Following quality control, sequencing data from 1772 nuclei from wild-type brains and 3968 nuclei from mutant brains were obtained. Uniform Manifold Approximation and Projection (UMAP) plots of these nuclei revealed a considerable diversity of wild type and mutant nuclei (Fig. 6A) Expression of a pan-mDA-neuron signature (*Th, Slc6a3, Nr4a2* and *En1)* was evident in a substantial proportion of the nuclei (Fig. 6B). To further characterize the snRNAseq data, distinct identities were assigned to different UMAP clusters based on the expression of markers previously described in the literature (Fig. 6C, Supplementary Fig. S4A)(*33*). Nuclei lacking robust expression of the pan-mDA neuronal signature, potentially isolated along with mCHERRY^+^ mDA neuron nuclei when sorting, were defined as astrocytes, oligodendrocytes (ODC) and non-mDA neurons based on typical neural cell markers. Based on the classification reported in reference 33, the six clusters which robustly expressed the mDA-neuronal signature were divided into: VTA1 (*Calb1^+^*/*Otx2^+^*), VTA2 (*Calb1^+^*/*Otx2^-^*), VTA3 (*Gad2^+^*/*Otx2^-^*), SNpc/VTA (*Sox6^+^*/*Aldh1a^-^*), SNpc-WT (>98.5% WT) and SNpc-KO (>98.6% KO) both groups *Sox6^+^*/*Aldh1a^+^* (Fig. 6C). Interestingly, one mDA-neuron groups (SNpc-WT) was almost exclusively enriched for wild-type nuclei (>98.5%) while another group (SNpc-KO) was almost exclusively enriched (>98.6%) for nuclei from mutant mice. In contrast, wild-type and knockout nuclei were distributed in roughly equal proportions in all other groups (Fig. 6A). SNpc-KO nuclei had diminished mDA neuron marker gene expression but did express *Sox6* and *Aldh1a1*, consistent with a relationship to SNpc mDA neurons. The expression of the SNpc markers *Sox6*, Aldh*1a1* and the VTA marker *Calb1* was distributed as shown in Supplementary Fig. S4B-C. Thus, these observations indicate a rather drastic influence on gene expression in SNpc neurons as compared to the other mDA neuron groups as a consequence of disrupted PRC2 function.

**Figure 6.**
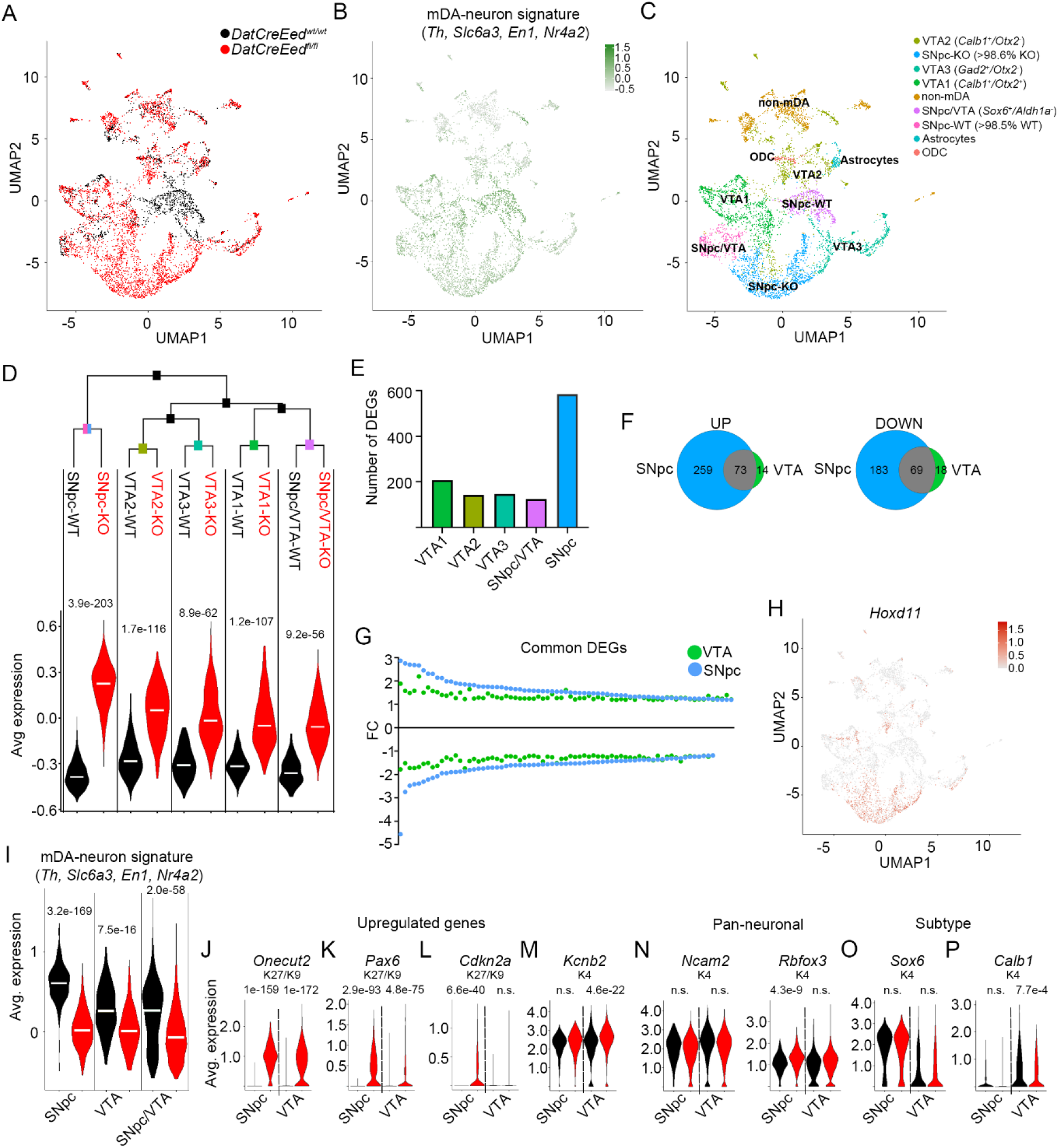
mDA neurons of the SNpc exhibit selective increased vulnerability to loss of PRC2 activity. (**A**). UMAP showing distribution of wild type (black) and mutant (red) nuclei. (**B**). Average expression levels of mDA-identity signature (*Th, Slc6a3, Nr4a2* and *En1*) in sequenced nuclei. (**C**). Classification of defined subgroups in sequenced nuclei, in UMAP space. (**D**). Hierarchical clustering of the mDA- neuron subgroups defined in (**C**) based on expression of 2000 most variable genes and composite expression score of the 25 most upregulated genes in the KO, plotted for the indicated groups. Black text denotes WT-nuclei and red text denotes KO-nuclei. (**E**). Number of differentially expressed genes (DEGs) in *DatCreEed^fl/fl^* mCHERRY^+^ single nuclei of the different mDA-neuron groups according to Fig. 6C. (**F**). Venn diagram of upregulated (UP) and downregulated (DOWN) genes in *DatCreEed^fl/fl^* mCHERRY^+^ single nuclei in combined VTA and SNpc. (**G**). Fold-change of gene expression (FC) for the DEGs common between VTA and SNpc for mutant vs. wild type. To put up- and downregulated genes on similar scales, the inverted fold-change (−1/FC) is plotted for downregulated genes. (**H**). UMAP visualization of *Hoxd11* expression in sequenced nuclei. (**I**). Average expression of mDA-neuronal signature in wild type (black) and mutant (red) nuclei in groups defined in (**C**), with VTA representing VTA1+VTA2+VTA3. (**J-M**). Violin plots exemplifying genes upregulated in both mutant SNpc and VTA (**J**), of genes more upregulated in mutant SNpc than in mutant VTA (**K**), of genes only upregulated in mutant SNpc and not in the VTA (**L**) of genes only upregulated in mutant VTA and not in SNpc (**M**). (**N- P**). Violin plots of pan-neuronal markers in wild type and mutant SNpc and VTA nuclei (**N**), of the SNpc- marker *Sox6* in wild type and mutant SNpc and VTA nuclei (**O**), of the VTA-marker *Calb1* in wild type and mutant SNpc and VTA nuclei (**P**). Wilcoxon Rank Sum test, with Bonferroni corrections for adjusted p-values. Adjusted p-values are included in panels **D** and **I-P**.

Hierarchical clustering based on the 2000 most variable genes in all nuclei from the mDA neuron groups revealed that the SNpc-WT and SNpc-KO clusters grouped separately from the VTA1-3 and SNpc/VTA groups (Fig. 6D, Supplementary Fig. S4D). To understand whether loss of PRC2-mediated repression had selective effects in different mDA-neuron subgroups, the signature expression of the 25 most upregulated genes in mutant nuclei were visualized in violin plots. This analysis revealed that the most profound increase occurred in the SNpc-KO vs. SNpc-WT groups (Fig. 6D, Supplementary Fig. S4E). We then generated a heatmap of all mDA-neuron groups based on the 205 differentially expressed transcripts between all mutant and wild type nuclei, of which 93 exhibited increased expression and 112 decreased expression. This revealed a block of genes that are strongly upregulated in the majority of SNpc-KO nuclei but only in subsets of nuclei in the VTA1-KO, VTA2-KO, VTA3-KO and SNpc/VTA-KO nuclei (Supplementary Fig. S4F). This group of genes was strongly enriched for PRC2 targets and for genes also upregulated in the bulk RNA-seq at 8 months.

To further understand the differential effects resulting from loss of PRC2 function, genes differentially expressed between wild type and mutant nuclei were analysed for each group separately. As expected, major effects on gene expression were only seen in mDA-neuron clusters derived from cells in which PRC2 had been disrupted by targeted knockout of *Eed* while marginal effects were seen in non-mDA-neuron groups. Notably, the largest number of DEGs was seen in the SNpc group (Fig. 6E) in which 584 genes were differentially expressed when comparing SNpc-KO vs SNpc-WT. In contrast, only 174 genes were differentially expressed when comparing wild type and mutant nuclei in the three VTA groups combined into one VTA group (Fig. 6F). The more substantial response to loss of PRC2 activity in the SNpc group was also reflected in the magnitude of increase or decrease in expression in common up- or downregulated genes between SNpc and VTA (Fig. 6G) and is also clearly illustrated by plotting highly upregulated genes, such as *Hoxd11,* in the UMAP plot (Fig. 6H). Moreover, expression of the mDA-neuron signature was more significantly reduced in the KO-nuclei of the SNpc group (Fig. 6I). Taken together, the greatest impact of *Eed* deletion occurred in the SNpc population which was also reflected by how TH immunoreactivity was substantially reduced in the SNpc at 8 and 16 months but was largely intact in the VTA (Fig. 3C-L).

Comparison of DEGs between mutant and wild type nuclei in the combined VTA and the SNpc showed equal (Fig. 6J, Supplementary Fig. S4G), stronger (Fig. 6K, Supplementary Fig. S4H) or exclusive (Fig. 6I, Supplementary Fig. 4I) upregulation of PRC2-targets in the SNpc. The few genes that were more robustly induced in the VTA are not PRC2 targets. Instead, they are typically expressed at robust levels in wild type cells and harbour the H3K4me3 modification (Fig. 6M, Supplementary Fig. S4J). Expression of pan-neuronal genes was not decreased in any of the groups, *Rbfox3* was actually slightly increased in the SNpc-KO nuclei (Fig. 6N). Notably, expression of the SN subtype specific gene *Sox6* was not decreased in the SNpc-WT vs SNpc-KO subgroups whereas expression of the VTA specific *Calb1* gene was reduced in the mutant VTA cells (Fig. 6O, P). We also plotted the expression of the four genes that constituted the mDA-neuron signature utilized Fig. 6B and F (Supplementary Fig. S4K).

To understand whether genes with specific chromatin states were regulated equally between mutant and wild type nuclei in VTA and SNpc, we utilized the chromatin states generated from the bulk ChIP-seq. As in the bulk RNA-seq, upregulated genes both in the VTA group and the SNpc group, were strongly enriched for the K9/K27 chromatin state (VTA: 8.3x increase over expected, p<4.2e-12, SNpc: 4.3x increase over expected, p<1.4e-13, Fisher’s exact test). In contrast, in the SNpc-group there was no significant enrichment of K4/K27 genes and in the VTA group the enrichment was less (2.9x increase over expected, p<0.0015, Fisher’s exact test) than for the K9/K27 state (Supplementary fig. S4L).

## Discussion

How long-term maintenance of cellular identity is coupled to permanent silencing of alternative lineages is largely unknown. This question is of particular interest for CNS neurons since their functional integrity and identity need to be maintained for several decades in the human brain. Our study reveals that in two well characterized neuronal populations of high clinical relevance, intact PRC2 function is essential for repression of aberrant gene expression as well as for the maintenance of cell type specific gene expression, but not for neuronal survial. Even though PRC2-mediated gene silencing has previously been shown to be required for proper neurogenesis in the neocortex (*7, 34*), the role of PRC2 in differentiated post-mitotic cells, such as neurons, is not understood. A previous report showed that in MSN and Purkinje cells, PRC2 is required to maintain silencing of death promoting genes (*6*). Similar “death-promoting” genes were upregulated in mutant mDA and 5HT neurons upon loss of PRC2 activity. However, there was no reduction in cell numbers, neither in mDA nor in 5HT neurons, even across an extended timespan (up to 16 months). Instead, there was a profound reduction in expression of subclass specific genes in both types of neurons. This is in contrast to a study wherein the methyltransferase *Ezh2*, another member of the PRC2 complex, was deleted in post-mitotic mDA neurons (*35*). Notably, deletion of *Ezh2* resulted in a selective and progressive loss of VTA neurons. Since the levels of K27 were not changed in the mutants, it is possible that this effect was uncoupled from the canonical methyltransferase function of *Ezh2*. In addition, in differentiated mDA neurons the expression level of *Ezh2* is lower than that of *Ezh1*, which also harbours methyltransferase capacity and thus could act as a redundant factor.

Since a majority of downregulated genes are not H3K27me3^+^ in the wild type cells and the main function of PRC2 is to maintain repression, the reduced expression is most likely an indirect effect of *Eed*-deletion. A similar effect was reported in MSNs and in differentiated β-cells wherein cell type specific genes were downregulated upon loss of PRC2-activity (*6, 36*). Utilizing the single-nuclei data set, we have tried to identify crucial upregulated factors that repeatedly correspond with a decrease in mDA-neuronal identity genes. However, we could not couple any single upregulated factor with the decrease in mDA identity genes. This would argue for a combined effect of several upregulated factors, which, during development harbour the capacity to induce other cell lineages as well as to silence the mDA or 5HT neuronal lineages.

Since neurons are post-mitotic, the progressive reduction of K27 in mutant neurons is not the result of failure to establish novel K27 marks during cell division. Previous reports that histone turnover in the rodent brain occurs at an extended time scale (∼220 days) (*37*) have recently been contrasted by studies showing that the histone variant H3.3 exhibit rapid and continuous turnover in differentiated CNS-neurons (*38*). Thus, the loss of K27 evident in the mDA and 5HT neurons likely is a consequence of combination of histone turnover and demethylase activity by *Kdm6a* and *Kdm6b,* both of which are expressed in mDA and 5HTneurons.

It has previously been proposed that the primary role for PRC2 in differentiated cells is to suppress transcription of bivalent K4/K27 genes (*6, 39*). Indeed, in both mDA and 5HT neurons there was an enrichment of K4/K27 bivalent genes among upregulated transcripts. However, presence of K9/K27 was associated with a higher probability of de-repression as well as a higher increase in both relative and absolute expression levels. This preference for activating K9/K27 genes over K4/K27 genes poses several questions. Is this a cell specific feature? What does it reveal about potential interactions between K9 and K27 associated factors, as well as how these modifications are interpreted by the transcriptional machinery? GO analysis of K9/K27 genes in 8month *DatCreEed^wt/wt^* mCHERRY^+^ nuclei clearly shows strong enrichment of categories such as regulation of transcription and early developmental categories, for example anterior/posterior pattern specification. K4/K27 genes on the other hand are more enriched for categories such as extracellular matrix organization, proliferation control and neuronal differentiation (Supplementary Fig. S1E). Notably, the categories enriched for the K4/K27 state reflects transitions from K4 state in neural progenitor cells (NPC) to K4/K27 state in mDA neurons as determined in our previous study (*20*). Similarly, the categories enriched for the K9/K27 state is reminiscent of silent genes carrying K27 already in NPCs but which gain K9 to become K9/K27 in mDA neurons. Thus, after terminal differentiation K9 appears to be gained as an additional layer of repression, hence the higher probability of de-repression of K9/K27 genes is an unexpected result. Especially, since in differentiated MSNs loss of PRC2 activity led to activation of predominantly poised K4/K27 genes (*6*). However, in that study, no analysis of K9 was performed, inactivation of PRC2 was achieved by targeting other components of the complex and a different Cre-promotor was used. Hence, differences could be due to different cell types, or the system used to delete PRC2 activity, which makes any direct comparison difficult. Interestingly, a recent study questions the whole concept of bivalency implicating that the combined presence of K4 and K27 does not represent a poised state (*40*). Indeed, our data showing that the K9/K27 state is a better predictor of de-repression than K4/K27 would reflect that K4/K27 does not represent a poised state wherein activation of transcription has a high probability to occur after loss of K27. However, further studies are required to solve this question.

De-repression of K9/K27-genes results in a more substantial absolute increase in expression when compared to K4/K27-genes. Given that the K9/K27-genes have nearly undetectable expression levels and the K4/K27-genes have substantially higher expression levels in the wild type cells the identified K9/K27 likely represent true K9/K27 promoters. The loss of K9 precedes the upregulation of expression, which implies that the loss of K9 is not a mere consequence of increased expression upon loss of K27. Rather, this suggests that presence of K9 at K27^+^ promoters is coupled to intact PRC2-function, alternatively intact K27 distribution. How loss of PRC2 activity and/or K27 promotes loss of K9 is not clear. It has previously been shown that PRC2 and K27 cooperate with K9 to maintain the K9 associated heterochromatin protein 1α (*41*). A more direct link between K27 and K9 has been reported for telomeric heterochromatin assembly, wherein PRC2 and K27 are essential for K9 as well (*42*). In differentiated mouse embryonic stem cells K9 is dependent on intact SUZ12 function (*43*).

The selective vulnerability to loss of PRC2 activity in mDA neurons of the SNpc is reminiscent of how the same cells are hypersensitive in the response to cellular stressors such as 6-OHDA. Thus, loss of identity upon deletion of *Eed* is mirrored by cell death upon increased cellular stress. Both vulnerability to cellular stressors as well as mouse genetic models of PD have been coupled to diverse processes such as mitochondrial dysfunction, inflammation, and protein misfolding, whereas the phenotype we report here is the consequence of dysregulation of transcriptional processes. Notably, the electrophysiological alterations of the *DatCreEed^fl/fl^* mutants are reminiscent of age dependent decline of similar parameters in the MitoPark mouse (*44*), whereas progressive loss of TH in the SNpc and progressive development of motor deficits mirror key aspects of PD. Thus, the more significant impact of loss of PRC2 function in SNpc mDA neurons shows that these cells harbour an additional selective vulnerability in addition to the death promoting effects of cellular stressors, inflammation and α- synuclein overexpression previously described. Whether this dual vulnerability is mechanistically coupled, remains to be addressed.

Expression of the homeobox gene *Engrailed1* (*En1*), which is a key survival factor for mDA neurons (*45*), is reduced in both the mutant SNpc and VTA (Supplementary Fig S4K). It has previously been showed that there is a pronounced reduction of K27 in mDA neurons of *En^+/-^* mice (*17*). Furthermore, in the same study *En1^+/-^* mutant exhibited heightened sensitivity to 6-OHDA treatment and reduced expression of both *Ezh1* and *Ezh2*. Hence, it appears that there is a link between *En1* and levels of K27, wherein En1 facilitates expression of *Ezh1* and *Ezh2,* thus helping to maintain K27 levels. Reciprocally, inhibition of PRC2 function in turn potentially leads to the upregulation of factors capable of repressing expression of *En1*. Given the fundamental role for En1 in the maintenance of mDA neurons it is possible that the loss of mDA neuronal traits is closely coupled to the reduced levels of *En1* in the mutants.

Taken together, our study elucidates how an epigenetic mechanism controls permanent gene silencing and maintenance of serotonergic and dopaminergic identity. It also reveals how loss of such epigenetic control does not compromise neuronal survival but leads to loss of subtype-specific function and to phenotypes that recapitulates symptoms characteristic of PD and mood disorders, providing a deeper understanding of how epigenetic mechanisms could contribute to the aetiology of these multifactorial diseases.

## Acknowledgements

Support was provided by The Knut and Alice Wallenberg Foundation (grant 2013.0075 to J.H., T.P., P.S. and K.C.), The Swedish Research Council (VR 2016-02536 to J.H.; VR 2016-02506 and VR 2020- 00884 to T.P.), The Swedish Brain Foundation (to J.H. and T.P.) and Torsten Söderbergs Stiftelse (T.P.). M.R. was financially supported by the Knut and Alice Wallenberg Foundation as part of the National Bioinformatics Infrastructure Sweden at SciLifeLab. The computations were enabled by resources in project SNIC 2021/23-184 provided by the Swedish National Infrastructure for Computing (SNIC) at UPPMAX, partially funded by the Swedish Research Council through grant agreement no. 2018-05973.

## Data Availability

The data sets generated and analyzed during the current study are available in the GEO repository with accession number GSE189018.

## Materials and Methods

### Ethical considerations

All animal experiments were performed according to Swedish guidelines and regulations, the ethical permits N189/15 and 6259-2020 was granted by “Stockholms Norra djurförsöksetiska nämnd, Sweden”.

### Mice

The generation of *DatCre, SertCre* (*23, 29*)*, Rpl10α-mCherry*(*46*) and *Eed^flo^* (*22*) mice has been previously described. Mouse lines were crossed and generated the *DatCreEed^fl/fl^Rpl10α-mCherry* and the *SertCreEed^fl/fl^Rpl10α-mCherry* lines used in our study. Mice were kept in ventilated cages with controlled 12 h light/dark cycles, temperature and humidity with water and food provided ad libitum. Mice were housed at a maximum number of four males or six females per cage. Both genders were represented in similar numbers for different type of experiments.

### Histological analyses

Animals were deeply anesthetized with Avertin intraperitoneal sodium pentobarbital (Apoteksbolaget AB) and perfused with room-temperature phosphate buffer saline through the ascending aorta, followed by ice-cold 4% paraformaldehyde. The brains were subsequently removed, postfixed in the same fixative for 16-18 h and cryoprotected for 24-48 h in 30% sucrose at 4°C, before being cut on a Leica microtome at 30 μm thickness. Sections were permeabilized in 5% BSA in PBS-Tx100 (PBS with 0.5% Triton-X100), followed by primary antibody incubation at 4°C for 16-18 h using sheep anti-TH (1:1000, cat# P60101-150, Pel-Freeze), anti-TPH2 (1:500, cat# T0678, Sigma), anti-H3K27me3 (1:500, cat# 9733, CST), anti-EED (1:500, cat# 85322, CST). Fluorescent detection was done with an Alexa-tagged secondary antibody from Molecular Probes, donkey anti-sheep (1:500, cat# A21448), goat anti-mouse (1:500, cat# A21151), donkey anti-rabbit (1:500, cat# A21206). Section images were obtained in confocal microscope LSM-700 from Zeiss

### Tissue processing for imaging and cell counting

Mouse brains were cleared using the CUBIC protocol (*47*) with minor modifications (*48*). In brief, mice were perfused with 4% paraformaldehyde and after post fixation, brains were washed in phosphate buffer (PB 0.1M, pH7.6-7.8) at 4°C for 24h. Brains were cut in 1mm slices using a brain matrix. 1 mm slices were incubated in CUBIC reagent 1 (25% urea, 25% N,N,N′,N′-tetrakis-(2- hydroxypropyl)ethylenediamine and 15% Triton X-100) at 37 °C for 2 days. Slices were transferred to fresh Cubic reagent 1 and incubated for further 24h at 37°C, before washing in PB (0.1 M) for 8 h at room temperature (8 x 1 h shaking). Tissue was incubated in blocking solution (5% BSA) for 24 h at 37°C and switched to anti-MCHERRY (1:5000, cat# AB0040-200, SICGEN) for 2 days at 37°C. After washing in PB (0.1 M) for 8 h at room temperature (8x 1h shaking), slices were incubated in Alexa 555, donkey anti-goat (1:500, cat# A21432, Invitrogen) for 24 h at 37°C and washed for 8h (8 x 1 h shaking) with PB (0.1 M) at room temperature. Slices were afterwards incubated in Cubic reagent 2 (50% sucrose, 25% urea, 10% 2,2′,2″-nitrilotriethanol, and 0.1% Triton X-100) while shaking for 16-18 h at 37°C. Tissue slices were placed in 1mm height chambers on a glass slide and imaged in Cubic reagent 2 in confocal microscope LSM-700 from Zeiss. Acquired images were analyzed with Imaris Cell Imaging software and cell bodies of mCHERRY positive cells were counted.

### FACS sorting of cell-type-specific nuclei

*DatCreEed^fl/fl^-Rpl10α-mCherry* and *SertCreEed^fl/fl^-Rpl10α-mCherry* mice were sacrificed with CO_2_ and brains were rapidly removed and transferred into cold PBS. The midbrain and hindbrain respectively, were dissected under fluorescent stereoscope and snap-frozen in dry ice. Tissue was thawed and dissociated using a 1 ml dounce homogenizer (Wheaton) in ice-cold lysis buffer (0.32M sucrose, 5nM CaCl_2,_ 3 mM MgAc, 0.1 mM Na_2_EDTA, 10 mM Tris-HCl, pH 8.0, 1 mM DTT, 1x complete proteinase inhibitor, EDTA-free (Roche). The homogenate was centrifuged for 5min at 500 x *g* and the pelleted nuclei were resuspended in a nuclear storage buffer (15% sucrose, 10 mM Tris_HCl, pH 7.2, 70 mM KCl, 2 mM MgCl_2_) supplemented with RNase inhibitor (RNase out, Invitrogen) and Proteinase inhibitor (Complete, Roche). When nuclei were used for 10x Genomics single cell RNA-seq, pelleted nuclei were instead resuspended in 2% BSA with RNase inhibitor. Re-suspended nuclei were filtered through a 30μm cup falcon (BD Biosciences, 340625) into a BSA-coated tube for sorting. Nuclei were stained with DAPI for 30 min prior to sorting.

FACS was performed using FACSAria Fusion cell sorter and the FACSDiva software (BD Biosciences). The nuclei were identified by forward- and side- scatter gating, a 561 nm laser with a 610/20 filter and a 405 nm laser with a 450/50 filter, quantifying DNA content per event to assure that only singlets were collected. A 100 μm nozzle, sheath pressure of 20-25 psi, and an acquisition rate of up to 1000 events per second were used. Cell type-specific nuclei were collected in batches of 1000, supplemented with a nuclear pellet buffer (10 mM Tris (pH 8.0), 100 mM NaCl, 2 mM MgCl_2_, 0.3 M sucrose, and 0.25% IGEPAL CA-630) to a volume of 10μl when used for bulk RNA-seq or ChIP-seq, whereas for 10x Genomics single cell RNA-seq, nuclei were preserved in 2% BSA until downstream procedure.

### Bulk RNA-sequencing and analysis

Sequencing libraries were generated from total RNA of 1000 FACS sorted cell-type-specific nuclei as previously described (*49*). In brief, total RNA was extracted from nuclei and the Smartseq2 protocol (*50*) was implemented to generate libraries. Sequencing was performed on an Illumina NovaSeq 6000 within the National Genomics Infrastructure in Scilife lab, Stockholm, Sweden. Raw 51 bp paired-end reads were aligned to the mouse genome (mm10 assembly) using STAR v2.7.0a with default settings (*51*). Gene expression was calculated using RPKM for genes (*52*). Differential gene expression analysis was performed using DESeq2 (*53*) with two separate DESeqDataSets, one for all mDA samples and one for all 5HT samples . A total of 35 mDA samples (9 wt and 10 mutant at 4 months, 8 wt and 8 mutant at 8 months) and a total of 16 5HT samples (4 wt and 4 mutant at both 4 months and 8 months) were used. Each dataset was filtered for genes with a least a total count of 10 summed across all samples. Differentially expressed genes were identified by requiring adj p<0.05 and using design formulas controlling for sex and contrasting wild type with mutant samples separately for 4 months and 8 months. GO-terms were obtained from Enrichr (*24, 25*).

### ChIP-sequencing and analysis

ChIP and library preparation for sequencing were performed as previously described in the ULI-NChIP protocol (*54*) with previously described minor modifications (*49*). Chromatin was IPed for 16-18h at 4°C with 0.25μg of anti-H3K27me3 (Cell Signaling,9733), anti-H3K9me3 (Active Motif,39161) or anti- H3K4me3 (Cell Signaling,9751). Libraries were sequenced on an Illumina NovaSeq 6000, 51 bp paired- end read.

A total of 64 ChIP-seq libraries were sequenced (two genotypes: wt and mutants, two time-points: 4 months and 8 months, four IPs: input, H3K4me3, H3K27me3 and H3K9me3, and finally four biological replicates for each combination). Reads were mapped to the mm10 mouse genome using bowtie2 v2.3.5.1 with default settings (*55*). Duplicate reads were marked using Picard v2.10.3. Coverage of mapped ChIP-seq libraries was generated using the tool bamCoverage in deepTools v3.1 with parameters ignoreDuplicates, binSize 50 and normalizeUsing RPKM (*56*). Signal-to-background relationships were investigated using the plotFingerprint tool in deepTools. Based on manual inspection of fingerprint and coverage plots, we decided to use all 64 samples in further analyses. The median fraction of duplicated reads for all samples was 34% (range 17–52). The median fraction of unmapped reads was 18% (range 7–60). The median total number of million mapped unique reads (after removal of duplicates) were 70 for inputs (range 38–78), 31 for H3K4me3 (range 17–40), 24 for H3K27me3 (range 8–63), and 60 for H3K9me3 (range 41–86).

For each histone modification and genotype, we identified marked genes by comparing the ChIP experiments to the input experiments using the csaw package version 1.22.1 in R (*57*) as previously described(*49*) with the following changes. Reads mapping to regions in the curated blacklist of problematic regions available as the bed-file ENCFF547MET from the ENCODE project were removed. Windows displaying an enrichment of reads above the global background were kept by requiring a minimal log2 fold-change (lfc) of two for all experiments except lfc=3 for K4 at 4 months and lfc=1.5 for K9 at 8 months. In each comparison, 4 biological replicates were used both for the ChIP and the input experiments. Marked genes were mapped to gene expression data based on gene symbols.

For visualizations, biological replicates were pooled. ChIP-seq coverage files of pooled samples were generated in deepTools using the bamCoverage function as for individual replicates. From the pooled coverage files, ChIP-seq heatmaps and average profile heatmaps were generated with deepTools using the plotHeatmap function and the plotProfile function, respectively. Coverages were plotted using the Integrative Genomics Viewer (IGV) version 2.8.3.

Enrichment for a specific chromatin state in a gene set was investigated by comparing the number of genes with the chromatin state in the gene set to the number of such chromatin states in a background set based on all genes using 2 × 2 contingency tables and Fisher’s exact test.

### Analysis of neurotransmitters by high-performance liquid chromatography (HPLC)

HPLC with electrochemical detection (ECD) were based on previously published protocols (*58, 59*). Briefly, ice-cold 0.1 M perchloric acid (PCA) was added to tissue sample, 50 µL PCA per 10 mg of tissue. Samples were incubated on ice for 10 minutes, vortexed and centrifuged at 16000 x g for 10 minutes at 4°C. Resulting supernatants were filtered through 0.2 µm nylon membrane inserts and centrifuged at 4000 x g for 5 minutes. Eluents were immediately stored at -80°C and subjected to HPLC-ECD analysis within 1 week. Standard solutions of: L-norepinephrine hydrochloride (NE), (+/-)-Epinephrine hydrochloride (EPI), 3,4-dihydroxyphenylacetic acid (DOPAC), 3,4-Dihydroxy-L-phenylalanine (DOPA), dopamine hydrochloride (DA), 5-hydroxyindole-3-acetic acid (5-HIAA), homovanillic acid (HVA), serotonin hydrochloride (5-HT), 4-Hydroxy-3-methoxyphenylglycol hemipiperazinium salt (MHPG), DL-4-Hydroxy-3-methoxymandelic acid (VMA) and 3-Methoxytyramine hydrochloride (3-MT) were prepared in 0.1 M PCA to obtain final standard concentrations of 200, 100, 50, 10, 5, 2 and 1 ng/mL. Calibration curves were obtained with the Chromeleon software through linear regression of peak area versus concentration. The HPLC-ECD system used was a Dionex Ultimate 3000 series (Dionex, ThermoFisher Scientific, USA). Analyte separation was performed on a Dionex C18 reversed-phase MD-150 3.2 mm x 250 mm column (3 µm particle size). Column and analytical cell were kept at 30 °C. The mobile phase, which was pumped at a flow rate of 0.4 ml/min, consisted of 75 mM monobasic sodium phosphate, 2.2 mM 1-octanesulfonic acid (OSA) sodium salt, 100 µL/L triethylamine (TEA), 25 µM ethylene-diamine-tetra-acetic acid (EDTA) disodium salt and 10 % acetonitrile (v/v), pH 3.0 adjusted with 85% phosphoric acid. For detection of neurotransmitters and metabolites, the first and second analytical cells were set to -100 mV and +300 mV, respectively. Processed tissue samples were thawed on ice in the dark for about 1 h before analysis, placed in the autosampler and kept at 5°C before injection. Chromatograms were acquired with Dionex Chromeleon 7 software over an acquisition time of 55 minutes. Analytes concentrations in tissue samples were expressed as ng/mg of tissue.

### Electrophysiology of mDA neurons

Mice were perfused with aCSF containing (in mM): NaCl (126), KCl (2.5), NaH_2_PO_4_ (1.2), MgCl_2_ (1.3), CaCl_2_ (2.4), glucose (10) and NaHCO_3_ (26). Their brains were rapidly removed and coronal brain slices containing SN 200 μm thick, were prepared with a microslicer (VT 1000S, Leica Microsystem, Heppenheim, Germany) in oxygenated (95% O_2_ + 5% CO_2_) ice cold modified artificial cerebrospinal fluid (aCSF) containing (in mM): NaCl (15.9), KCl (2), NaH_2_PO_4_ (1), Sucrose (219.7), MgCl_2_ (5.2), CaCl_2_ (1.1), glucose (10) and NaHCO_3_ (26). Slices were incubated, for 1 h at 32 °C and thereafter at 28 °C, in oxygenated modified aCSF containing (in mM): NaCl (126), KCl (2.5), NaH_2_PO_4_ (1.2), MgCl_2_ (4.7), CaCl_2_ (1), glucose (9) and NaHCO_3_ (23.4). Slices were transferred to a recording chamber and were continuously perfused with oxygenated aCSF at 32-34 °C.

Whole-cell patch-clamp recordings of visually identified DA neurons in the SN were made as described previously (Yao et al., 2018). Borosilicate patch electrodes (3-5 MΩ) were filled with a solution containing, in mM: 120 D-gluconic acid potassium salt, 20 KCl, 2 MgCl_2,_ 1 CaCl_2,_ 10 HEPES, 10 EGTA, 2 MgATP, 0.3 Na_3_GTP, pH adjusted to 7.3 with KOH. Whole-cell membrane currents and potentials were recorded with a MultiClamp 700B (Axon Instruments, Foster City CA, USA), acquired at 10 kHz and filtered at 2 kHz. Data were acquired and analyzed with the pClamp 11 software (Axon Instruments, Foster City CA, USA).

### Viral tracing injections

Mice were anesthetized with 4mg/ml isoflurane supplemented in the air, while placed in a separate cage and afterwards they were mounted in a stereotaxic frame (David Kopf Instruments, Tujunga, CA) equipped with a mouse adapter. Mice were kept anesthetized throughout the procedure by inhaling isoflurane at 2mg/ml concentration. Anterograde tracing of the midbrain dopaminergic neurons was achieved by injecting a Cre-dependent adeno-associated virus (AAV) expressing EGFP (AAV2/2.pCAG.FLEX.EGFP.WPRE.bGH)(*26*) in the midbrain of 8 months old mice. Using Bregma as a reference point, 1μl of the AAV virus was injected unilaterally reaching the SN (anteroposterior (AP): -2.9, mediolateral (ML): -1.25, dorsoventral (DV): -4.5) and 1μl reaching the VTA (anteroposterior (AP): -3.1, mediolateral (ML): -0.5, dorsoventral (DV): -4.2). All coordinates are millimeters relative to Bregma according to *The Mouse Brain in Stereotaxic Coordinates (Academic Press, San Diego, CA, 2012).* Virus injection was executed at 0.2μl/min dispense rate. Animals were sacrificed 3 weeks after surgery by CO_2_.

## Behavioral Experiments

### Open field

For assessment of general ambulatory ability, mice were placed in 45 x 45 x 28 cm plastic, non-transparent boxes and activity was recorded for 60 min using the Ethovision XT 15, Noldus software. Activity is measured as mean of total distance covered in 1 hr with 5 min time bins.

### Pole test

Motor coordination and movement initiation were assessed by the pole test. To perform this task, mice were placed on the top of a wooden pole (50cm height, 1cm diameter) that is fixed to a wooden stable base. Each animal was placed with all four limbs grasping the pole and facing the tip of it. The total time they spent to turn themselves downwards and climb down the pole was calculated.

The day before the experiment, mice were trained to orient themselves and descent the pole for 5-10 times. On the test day, each animal was recorded performing the test for 5 times and the average time for every mouse is presented. In order to avoid exhaustion of the mice, a maximum time for every trial was defined at 120s, time score that was also given to objects that failed the experiment, by either not turning themselves downwards at all or by descended the pole by rolling down.

### Elevated Plus Maze

The Elevated Plus Maze (EPM) test evaluates anxiety-like behavior in mice. The set up consisted of a cross with two open arms and two enclosed ones, elevated ∼1 meter from the ground. Mice were placed in the centre of the cross, facing towards the open compartment. Mobility and preference over the two types of arms were recorded for 5 min using Ethovision XT, Noldus software.

### Rearing

Locomotion activity and exploratory behavior were evaluated by the rearing events. Mice were placed in a plexiglass 15 cm-diameter cylinder and videotaped for 10 minutes. Vertical rearing events supported on the cylinder wall or unsupported were manually counted. Mice were tested individually, without any visual interaction with each other.

### Forced Swim Test

The Forced Swim Test (FST) or Porsolt test (*60*) was used to evaluate depression-like traits in rodents. More specifically mice were forced to swim in a plexiglass cylinder (15cm-diameter 30cm height) halfway filled with water at 25±C for 6 min. During the session mice were recorded and time of immobility was assessed by a trained observer. The first two minutes were counted as habituation time and the left four minutes served as the actual experiment. During those minutes, as immobility time was counted the time mice spent floating in the water without any effort to move but only the necessary moves that would let them keep the head above the surface. Video recordings were scored twice from the same observer and the average of those scores were calculated.

### Rotarod

Motor coordination was evaluated by utilising the rotarod test with increasing speed. Mice were familiarized with the apparatus for three trials at a fixed speed of 4rpm. After a resting period of 2 hours, mice performed the test with increasing speed over time. The acceleration protocol spans from 4 rpm to 38 rpm within 5 min, where the latency to fall was measured. Every mouse participated 3 times and the average time was calculated for every individual.

### Grip Strength

Mice were placed horizontally on a metal net-shaped frame with all four limbs and instantly the apparatus was twisted 180°. With that set up, the back of the mice was facing the ground and the time they managed to hold gripped on the frame was calculated. Each individual was monitored for 3 trials and the average time was noted.

### Conditioned Place Preference

Experiment was performed in a rectangular apparatus with 3 chambers measuring 15×25 cm each. One compartment was coloured grey, the middle one was white and the last one was coloured with black and white stripes. One the first day mice were allowed to explore the whole apparatus for 20 min. On day 2 and 3 mice were confined to one compartment and injected with cocaine (20mg/kg) or saline. After injection they remained in the compartment for 30 min. On day 4 and 5 the same pairings were repeated. Pairing of drug and compartment was counterbalanced across animals. On day 6 mice were placed in the middle compartment and were freely allowed to move between all 3 of them. While there was no injection on that day, mice were recorded and their preference for every compartment was calculated.

The following day all mice were injected with cocaine (20mg/kg) and their motor skills and mobility were examined in the Open Field test, where they were recorded for 60 minutes.

### Single nucleus RNA sequencing and analysis

Tissue from mouse ventral midbrain (3 control and 3 KO) were used to obtain the single nuclei (1883 control & 4103 KO). Single nucleus libraries were made using Single Cell 3’ v3 on Chromium platform (10X Genomics & SciLifeLab, Stockholm) in accordance with the manufacturer’s protocols. Libraries were sequenced on a NovaSeq 6000 system (NGI, SciLifeLab, Stockholm).

Sequenced reads were demultiplexed and aligned to Transcriptome: mm10-3.0.0_premrna using Cell Ranger Pipeline Version 3.1.0 (10X Genomics). Quality control and filtering of data were performed in multiple steps. First, percent_mito% and percent_ribo% were computed based on the percentage of transcripts that map to mitochondrial genes and ribosomal genes respectively. Then, CellCycleScoring (Score cell cycle phases); S.Score, G2M.Score, and Phase columns were calculated based on the expression of G2/M and S phase canonical markers. Next, average relative expression of each gene per cell i.e., the ratio of a gene’s UMI counts to the sum of all UMIs per library (nucleus), was calculated. *Malat1* was on average the most abundant gene per library (nucleus), with a mean fraction of total UMI counts per library of about 2% and being expressed in all nuclei*. Malat1* is frequently detected in poly-A captured RNA-seq libraries, independent of the used protocol. However, compared to other methods it is even more abundant in snRNA-seq and therefore used to estimate the nuclear proportion of total mRNA (https://doi.org/10.1371/journal.pone.0209648). *Malat1* was filtered out for downstream analysis. Genes detected in fewer than 3 libraries were removed. Libraries with less than 500 detected genes, more than 10,000 detected genes (doublets) and with a percent mito> 0.9 were also filtered out. After the filtering steps, 1772 control and 3968 KO nuclei remained. The remaining nuclei have UMI/library over 620, gene/library over 500, with average UMI/library 23474 and average gene/library 4965.

Then a Seurat object was made with this filtered data (Seurat version ‘4.0.1’). Data was log-normalized with Seurat NormalizeData function, and a scale.factor of 10e^4^. Next, the 2000 of most highly variable genes (HVGs) were identified using the FindVariableFeatures function (selection.method = “vst”, with clip.max = (n.Cells)^2^). The log-normalized data were then scaled and centered using the ScaleData function (model.use = “linear”). Principal Component Analysis (PCA) was done using the 2000 HVGs, which reduced the dimensionality of the data into the calculated components whilst maintaining the most important gene expression differences across libraries.

We identified the most significant PCs based on the JackStraw procedure using the JackStraw (num.replicate = 100) and ScoreJackStraw functions. After plotting the JackStraw scores, noticeable gaps in p-values were observed at PCs 18, 22 and 32. We also used a heuristic method called ElbowPlot to visualize the standard deviations of the principal components and at PCs 18, 22, and 32 there were noticeable inflection points (“elbows”). We chose to continue with the first 32 PCs, which we think contain most of the variance without losing any true signal that reflects biological heterogeneity. Next we used PCs 1-32 as input to the Seurat FindNeighbors function (k.param = 20, dims = 1:32), as well as in the FindClusters function with the resolution parameter between 0.3 and 2.0 in 0.1 intervals, using the Louvain algorithm. We found the most optimal resolution for res = 0.3 with 18 clusters (a purely heuristic approach based on known markers for cell types, neuroanatomical regions and hierarchical dendrograms). The final 9 subgroups were created by combining related clusters together and based on markers in UMAP and hierarchical dendrograms. The same PCs 1-32 were used in the Seurat function RunUMAP. Subgroup-enriched markers (upregulated & downregulated) were identified using Wilcoxon Rank Sum test (logfc.threshold = 0.25, min.pct = 0.1, min.diff.pct = -Inf, only.pos = FALSE). In this function, p-values were adjusted for multiple comparisons with the Bonferroni correction.

DEG was also performed between all control nuclei and all KO nuclei, irrespective of clusters/ subgroups, using the FindMarkers function with the same setting and parameters as above.

Hierarchical clustering was done for all mDA subgroups (SNpc, VTA1-3, SNpc/VTA), by genotype (WT, KO), using the 2000 most highly variable genes and the Seurat BuildClusterTree function. This clustering results in a phylogenetic tree based on calculating the ’average’ cell from each mDA subgroup_genotype “identity class”. The distance matrix for this tree was calculated in gene expression space.

Heatmaps showing the clustering analyses, using either DEGs between control and KO (all nuclei irrespective of cluster) or the 2000 HVGs across the subgroups split by genotype (subgroup_WT, subgroup_KO) were generated using ComplexHeatmap version2.6.2(*61*).

### mDA-neuron signature (*Th, Slc6a3, En1, Nr4a2*)

A composite expression score of a mDA neuron gene set (*Th, Slc6a3, En1, Nr4a2*) was used. The Seurat AddModuleScore function was used to calculate the average expression levels of this gene set (signature) on single nuclei level. Genes in this gene set are binned based on their averaged expression, and the control features are randomly selected from each bin.

The same function was applied to calculate the signature score forthe top 25 most highly upregulated genes in the KO nuclei when compared to the WT nuclei, irrespective of their clusters.

