## Supplementary figures for "PRC2-mediated repression is essential to maintain identity and function of differentiated dopaminergic and serotonergic neurons"

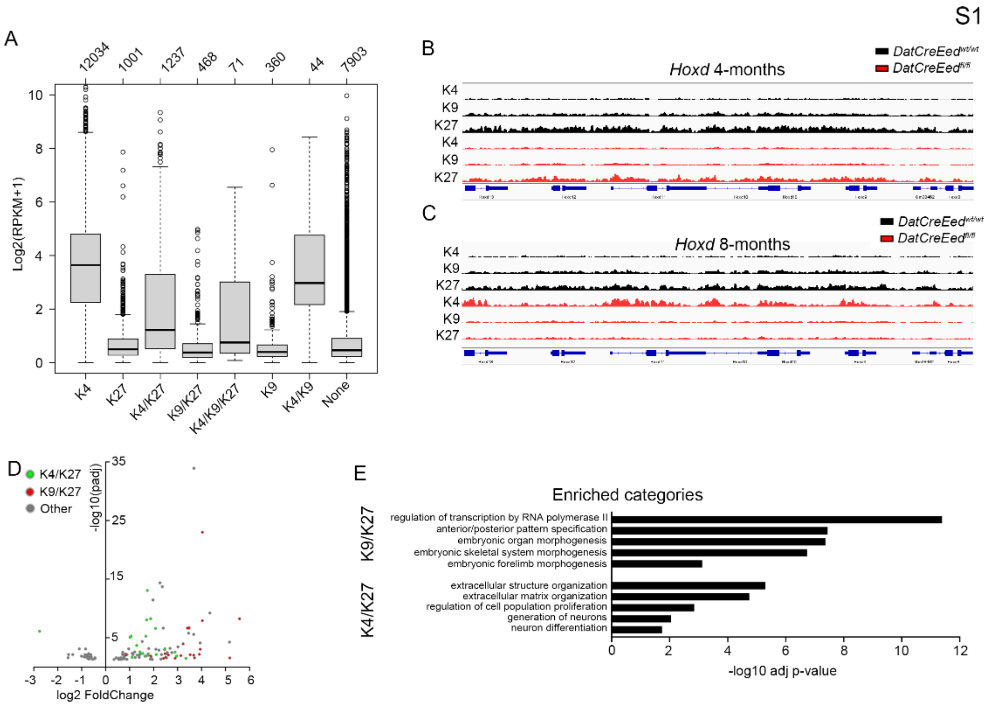


**S1. At 4-months, individual chromatin states are associated with different levels of gene expression.**

(**A**). Boxplots showing the expression levels (log2(RPKM + 1)) of genes in the different chromatin-state categories. The center line is the median, bounds are the 25th and 75th percentiles, and whiskers are ±1.5 IQR. Chromatin states with an average gene expression that is different compared to the global average gene expression are indicated by *p*-values obtained by a two-sided Wilcoxon rank-sum test. (**B**). IGV-tracks of K4, K9 and K27 at the *HoxD*-cluster at 4 months. (**C**). IGV-tracks of K4, K9 and K27 at the *HoxD*-cluster at 8 months. (**D**). Volcano plot showing differentially regulated genes at 4 months in isolated mCHERRY$^{+}$-nuclei from *DatCreEed*$^{fl/fl}$ ventral midbrain. Genes are labelled as belonging to H3K4me3/H3K27me3 (green) or H3K9me3/H3K27me3 (red) chromatin states in WT cells. (**E**). Enriched categories from “GO Biological Process” for genes with the K9/K27 and K4/K27 chromatin states.

S2

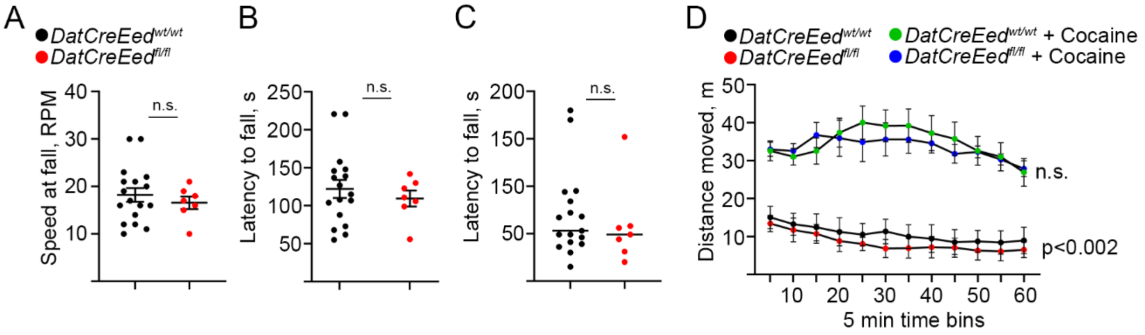


**S2** (**A**). No significant difference in speed of rotation at fall in rotarod test between *DatCreEed*$^{wt/wt}$ and *DatCreEed*$^{fl/fl}$ mice. (**B**). No significant difference in latency to fall in rotarod test between *DatCreEed*$^{wt/wt}$ and *DatCreEed*$^{fl/fl}$ mice. (**C**). No significant difference in latency to fall in grip strength test between *DatCreEed*$^{wt/wt}$ and *DatCreEed*$^{fl/fl}$ mice. (**D**). No significant difference in response to cocaine between *DatCreEed*$^{wt/wt}$ and *DatCreEed*$^{fl/fl}$ mice in the open field test. In **A-B**, significance was calculated with unpaired t-test. In **C**, p-values calculated by Two-way repeated measures ANOVA.

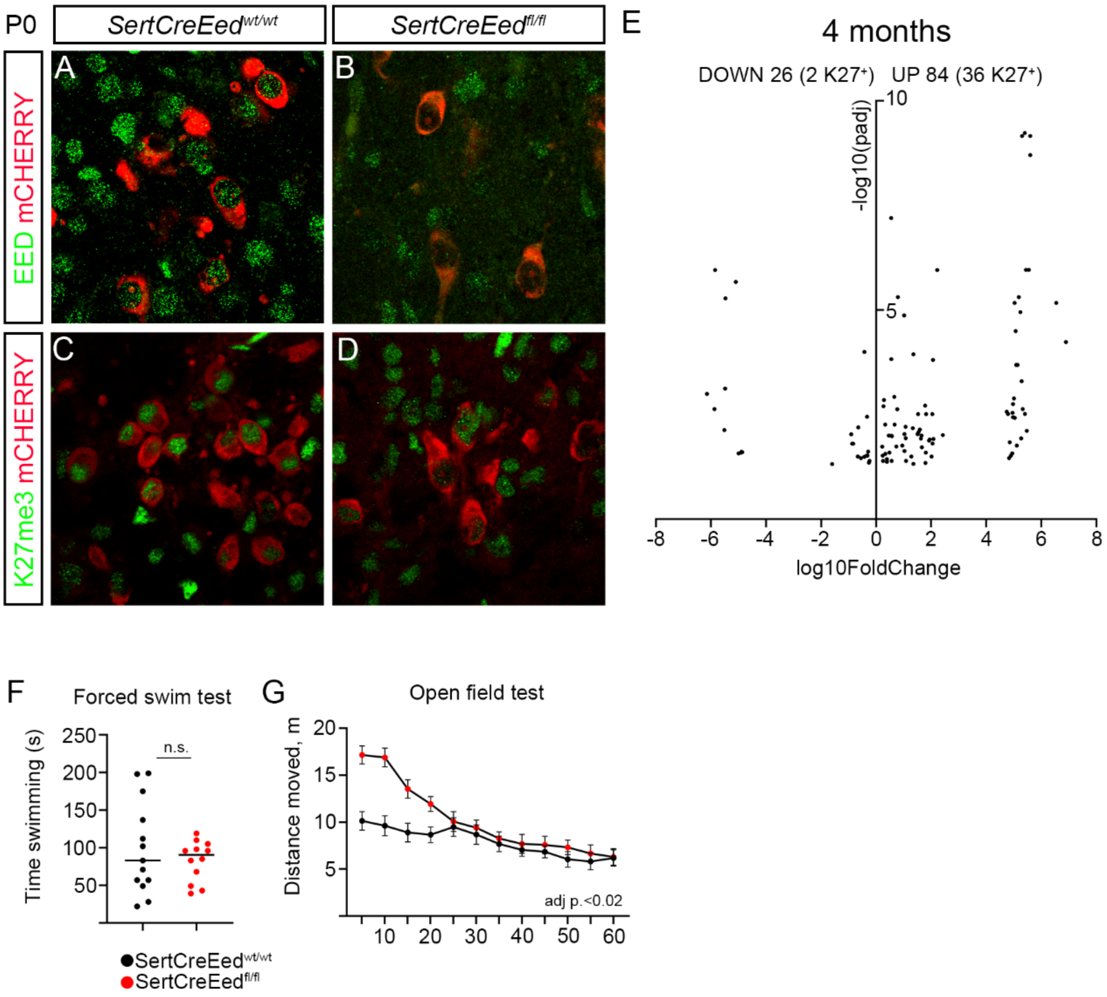


**S3.** (**A**). Overlap between EED and mCHERRY immunostaining at P0 in *SertCreEed*$^{wt/wt}$ dorsal raphe. (**B**). Loss of EED in mCHERRY$^{+}$-cells at P0 in *SertCreEed*$^{ft/fl}$ dorsal raphe. (**C**). Overlap between H3K27me3 and mCHERRY immunostaining at P0 in *SertCreEed*$^{wt/wt}$ dorsal raphe. (**D**). Overlap between H3K27me3 and mCHERRY immunostaining at P0 in *SertCreEed*$^{fl/fl}$ dorsal raphe. (**E**). Volcano plot showing differentially regulated genes at 4 months in isolated mCHERRY$^{+}$-nuclei from *SertCreEed*$^{fl/fl}$, the number of K27$^{+}$ genes are indicated within brackets. (**F**). No significant difference in time spent swimming in the forced swim test between *SertCreEed*$^{wt/wt}$ and *SertCreEed*$^{fl/fl}$ mice. (**G**). Initial difference in distance moved between *SertCreEed*$^{wt/wt}$ and *SertCreEed*$^{fl/fl}$ mice in open field test. In **F** significance calculated with unpaired t-test. In **G** p-value calculated by Two-way repeated measures ANOVA.

S4

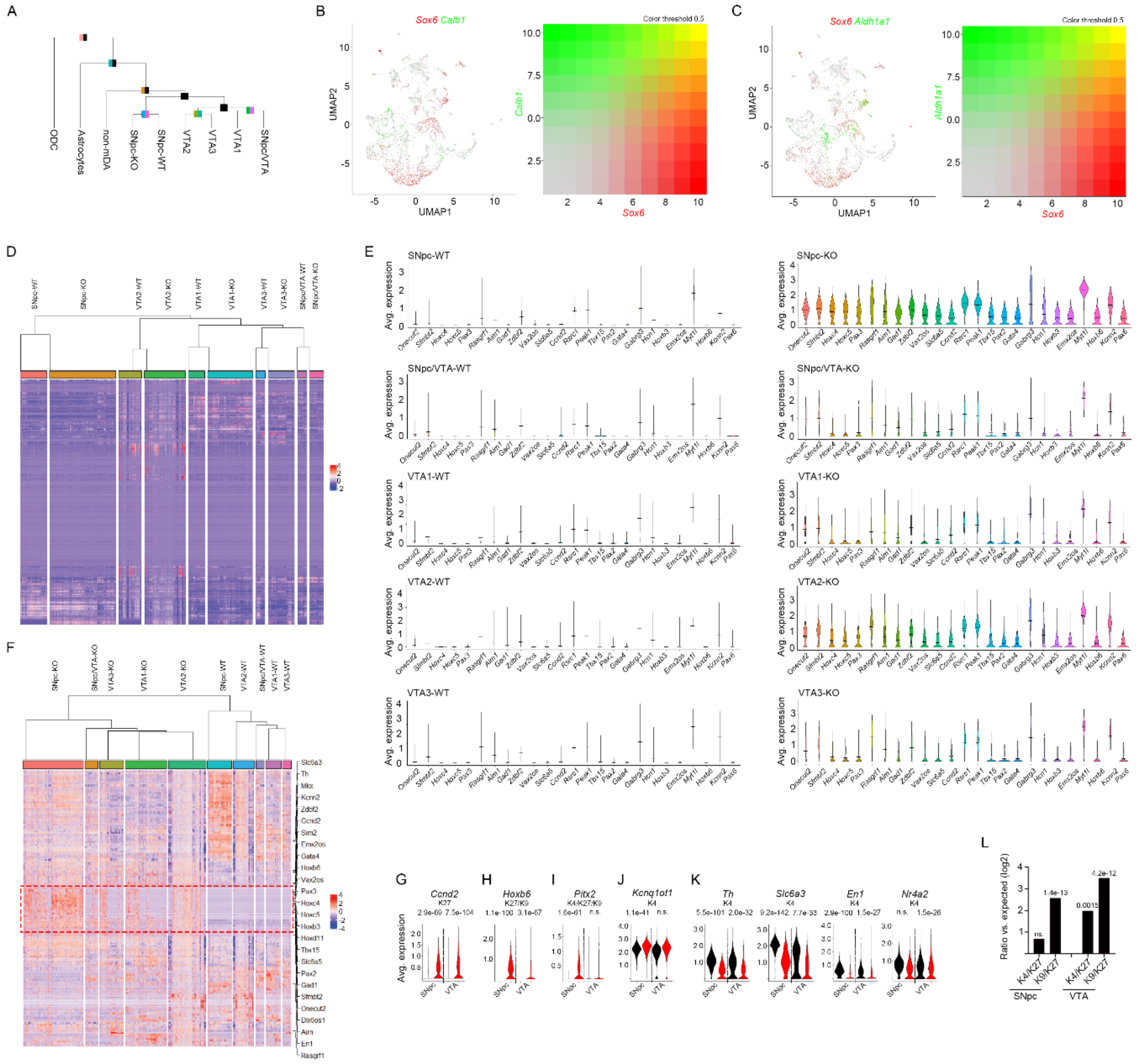


**S4.** (**A**). Hierarchical clustering of all nuclei, utilizing the 2000 most highly variable genes. (**B**). Expression of *Sox6* and *Calb1* in the sequenced mCHERRY$^{+}$-nuclei (**C**). Expression of *Sox6* and *Aldh1a1* in the sequenced mCHERRY$^{+}$-nuclei. (**D**). Hierarchical clustering and heatmap utilizing the 2000 most highly variable genes in mDA-nuclei. (**E**). Violin plots of the signature score for the 25 most upregulated genes in the mutant, plotted per mDA-neuron WT and KO groups. (**F**). Hierarchical clustering and heatmap utilizing the DEGs between m-DA-neuron KO and WT nuclei. (**G-J**). Violin plots exemplifying genes upregulated in both mutant SNpc and VTA (**G**), of genes more upregulated in mutant SNpc than in mutant VTA (**H**), of genes only upregulated in mutant SNpc and not in the VTA (**I**) of genes only upregulated in mutant VTA and not in SNpc (**J**). (**K**). Violin plot of the genes included in the mDAidentity profile in wild type and mutant SNpc and VTA nuclei. (**L**). Enrichment of K4/K27 and K9/K27 chromatin states for upregulated genes in *DatCreEed*$^{fl/fl}$ mCHERRY$^{+}$ single nuclei from SNpc and VTA. Wilcoxon Rank Sum test, with Bonferroni corrections for adjusted p-values (**G-K**). Fisher's exact test (**L**).
